# A generic multi-level stochastic modelling framework in computational epidemiology

**DOI:** 10.1101/491605

**Authors:** Sébastien Picault, Yu-Lin Huang, Vianney Sicard, Thierry Hoch, Elisabeta Vergu, François Beaudeau, Pauline Ezanno

## Abstract

There is currently an overwhelming increased interest in predictive biology and computational modelling. The development of reliable, reproducible and revisable simulation models in computational life sciences is often pointed out as a challenging issue. Population dynamics, including epidemiology, has not yet developed a language to formalize complex models in a univocal and automatable way, hence hindering the capability to implement in short time reliable, revisable and expert-friendly models intended for realistic mechanistic simulations. In epidemiology specifically, models aim not only at understanding pathogen spread but also at assessing control measures at several scales. To achieve this goal efficiently, best software practices should be supported by Artificial Intelligence methods to handle experts’ knowledge. The framework EMULSION presented here intends to both tackle multiple modelling paradigms in epidemiology and facilitate the automation of model design. We therefore built both a domain-specific language (DSL) for the modular description of complex epidemiological models, and a generic simulation engine designed to embed existing modelling paradigms within a homogeneous architecture based on adaptive software agents. The diversity of concerns (biology, economics, human activities) involved in real pathosystems requires an explicit, comprehensive and intelligible way to describe epidemiological models, to involve experts without computer science skills throughout the modelling, simulation and output analysis steps. This approach was applied to compare hypotheses in modelling a zoonosis (Q fever), to study its transmission dynamics within and between cattle herds at a regional scale, and to assess the contribution of transmission pathways. Separating model description from the simulation engine allowed epidemiologists to be involved in assumption revision, while guaranteeing very few code modifications. We assessed the added value of EMULSION by applying the DSL and the simulation engine to a concrete disease. Future extensions of EMULSION towards a broader range of epidemiological concerns will reduce significantly the time required to design and assess models and control measures against endemic and epidemic diseases. Ultimately, we believe this effort is a major lever to increase scientists’ preparedness to face emerging threats for public health and provide rapid, reliable, and reasoned assessments of control measures.

## Introduction

### Balancing development time, reliability and intelligibility in computational models

Computational modelling is essential to better explore complex systems. In particular, agricultural production systems present highly coupled biological, farming, environmental, and economic processes, involving a diversity of interacting entities, from individual scale up to whole territories. Their analytical investigation is strongly limited by the interplay between all processes and scales, leading to high dimensional and highly nonlinear systems, but also by the boundaries of knowledge concerning the exact interactions between actors of such systems. Mechanistic simulation models can assess the relevance of assumptions by comparing model outputs to field data, provide predictions on systems evolution under real or counterfactual scenarios, and help identify levers to control those systems. However, it is crucial that alternative hypotheses and practicable actions be tested in short time, in strong interaction with experts, to quickly identify assumptions providing the most significant insights, or actions driving the system to a desired state. Such “sieving” of hypotheses also promotes parsimonious models, highlighting key elements, hence allowing for deeper understanding and easier comparisons.

However, developing simulation codes directly from models requires strong skills in computer programming. Any change in hypotheses, scenarios, model structure or even just parameters is excessively time-consuming to foster incremental design of models and expert involvement. Also, reliability and reproducibility issues of ad-hoc simulation programs threaten conclusions drawn from computer experiments. To avoid misinterpretations coming from programming biases [1], several good practices in software development were proposed [2] (e.g. precise code documentation, systematic testing, versioning, etc.), but erroneous programs can also reach such standards [3].

Models are intended to change with biological knowledge and research questions. Assessing their relevance rather than simulation code quality requires to allow experts scrutinizing their very components (parameters, functions, modelling paradigms, contact structures, etc.), instead of their implementation within a specific programming language. It is then the responsibility of computer scientists to provide an automated, reliable and rapid translation into code. Our approach is in line with this mindset, by coupling a modular simulation architecture with a Domain-Specific Language (DSL), which gives experts the ability to understand and design the multiple components of an epidemiological model without programming.

### The diversity of modelling and related issues in epidemiology

Epidemiology is an epitome field for addressing such issues. Since Kermack and McKendrick’s seminal works [4], the complexity of models increased to allow for realistic decision support at several scales [5, 6], incorporating a broad range of concerns: infectious processes, demography, environmental conditions, underlying contact structure provided by transportation or trade, etc. Models became hard to design and harder to implement in a reliable way, because life scientists are not expected to master programming skills [7]. The diversity of modelling paradigms, as presented below, from rather formal and analytical, to rule-based descriptions of processes involved in the system, also reduces the ability to revise or compare models in response to evolving scientific knowledge and purposes. This often results in heterogeneous, ad-hoc simulation programs which cannot be compared, enhanced, even used, without diving into the code. However, responsiveness in modelling and in scenario assessment is a stake to provide relevant control measures against outbreaks, especially in the case of an animal health crisis.

Compartment-based models (CBM) [8] describe disease dynamics by state variables (amount or proportion of individuals in each health state). CBM can also represent demographic dynamics with input and output rates, and incorporate additional concerns (e.g. age structure, species, or environment-borne contamination) by splitting compartments. CBM assume that individuals differ only by a few discrete variables which determine their compartment. To assess targeted control measures, the multiplication of compartments and transitions required to account for finer-grained features can make the model very like individual-based models (IBM) [9]. These latter keep individual diversity explicit [10, 11] and represent them with their behaviors, environment, possible goals (e.g. [12–16]). This comprehensive understanding of causal mechanisms occurring in biological systems allows to compare individual trajectories and measure the impact of fine-grained actions [17]. The capability to increase indefinitely detail level as needed is counterbalanced by a high computational cost and by a difficulty to calibrate parameters (even with parsimonious models), two major drawbacks of IBM, since the scientific soundness of simulation outcomes relies upon repetitions and sensitivity analysis. Their use on very large scales (e.g. millions of agents) is a challenge, even with massively parallel platforms, strong software optimizations and oversimplified epidemiological assumptions [13, 18]. Metapopulations approaches [19, 20] have been applied in epidemiology for handling region-wide models at a moderate computational cost. Populations are modelled in interaction through a contact structure [21] representing neighborhood relations, transportation or trade exchanges [22, 23], or vector-or airborne transport processes. Yet, approximations in sub-populations dynamics may result in overestimating infections [24, 25] compared to equivalent IBM.

Most paradigms share the flow diagram formalism, with nodes denoting health states (possibly combined with other concerns), and transitions labeled with rates. In continuous, deterministic approaches, they are equivalent to an Ordinary Differential Equation (ODE) system, while in stochastic models, rates can be used, after conversion into probabilities, either in discrete event methods (e.g. the Gillespie algorithm [26]), or in multinomial sampling in discrete time approaches [27]. The main drawback of flow diagrams is that several concerns (infection, demography, herd management…) often are mixed together in a single representation, reducing the readability of the model, while other features (parameters, processes, data…) are not systematically explicitly depicted (e.g. the exponential distribution of state durations, or pathogen shedding during infectious states). Then, when writing actual simulation code, several implicit assumptions or actions are just added on the fly, which hinders early model comparison and often leads to biases in the late stage.

### Related computer science solutions and specificities of our approach

Epidemiology does not provide any systematic methodology for designing, implementing or assessing the diversity of its models yet. Other life sciences, which have long faced major computational problems, have adopted powerful formalisms to express their models in a quite explicit and comprehensive way and automatize their development. For instance, in molecular biology, the Systems Biology Graphical Notation (SBGN) [28] offers a visual syntax for describing reactions, compounds or feedback loops. In epidemiology, such attempts are still at their early stage. Formalisms inspired from multi-scale processes in physics [29], or proposing a strong complexification of flow diagrams [30], are not likely to facilitate the appropriation of models by epidemiologists. Conversely, the ODD protocol (“Overview, Design concepts, Details”, [31]) aims to obtain comprehensive knowledge from disciplinary experts within a textual template; however, feedbacks on its actual use for designing models emphasize ambiguities and “the lack of real specifications” [32].

Most simulation programs developed for implementing epidemiological models are hand-written ad-hoc tools dedicated to a single pathogen in an applicative context, to evaluate a specific set of control measures. Reliability of such codes inherently depends on the programming skills of their authors, and prove difficult to use and maintain in the long-term. Especially, instead of high-level programming languages (Scilab, R, Python…), performance considerations lead to using low-level ones (C++), yet harder to master, debug and maintain. Besides, even in object-oriented development, abstraction is rarely used (excepted e.g. in [33] on vector-borne disease mechanisms).

However, classical general-purpose simulation platforms tend progressively to be used, for instance GAMA [14], NetLogo [16] or Repast [34]. They provide indeed reliable, ready-made tools for calculations or data integration, leaving more time to focus on modelling itself. Yet, they are not specifically designed for epidemiology and still require a significant time spent on software development. Simulation libraries and platforms dedicated to epidemiological issues are rising, e.g. SimInf [35], a R library for data-driven CBM; MicroSim [36], an agent-based platform for several diseases; or GLEaMviz [37], a metapopulation-oriented platform. To our knowledge, the most advanced approach from a software engineering viewpoint is Broadwick [38], a framework for CBM and IBM with interaction networks, which still requires writing large portions of code to derive specific classes and carry out simulations on practical cases. Another interesting approach, though dedicated to a specific study, relies upon geographical levels and a separation of activities to define efficient aggregations of individuals [39]. KENDRICK [40] defines a DSL fostering a strong separation of concerns (infectious dynamics, spatial distribution, species), and generates C/C++ code to run simulations efficiently. But, it only targets CBM with theoretical (SIR-type) models.

The framework EMULSION (for “Epidemiological Multi-Level Simulation”) we propose is to our knowledge the only contribution that combines the capability of integrating several modelling paradigms and several scales with dynamic aggregation levels (through a multi-level multi-agent architecture to wrap them), and the explicit description of models (through a DSL dedicated to epidemiolocal issues) [41] (Fig 1). Our objective was indeed to 1) define a formalism making models as accurate as possible, so that a comprehensive description can be shared amongst experts, and its implementation automatized; and 2) encompass existing modelling paradigms within a common interface, to make them interchangeable, or even combine them as proposed by [42].

**Fig 1.**
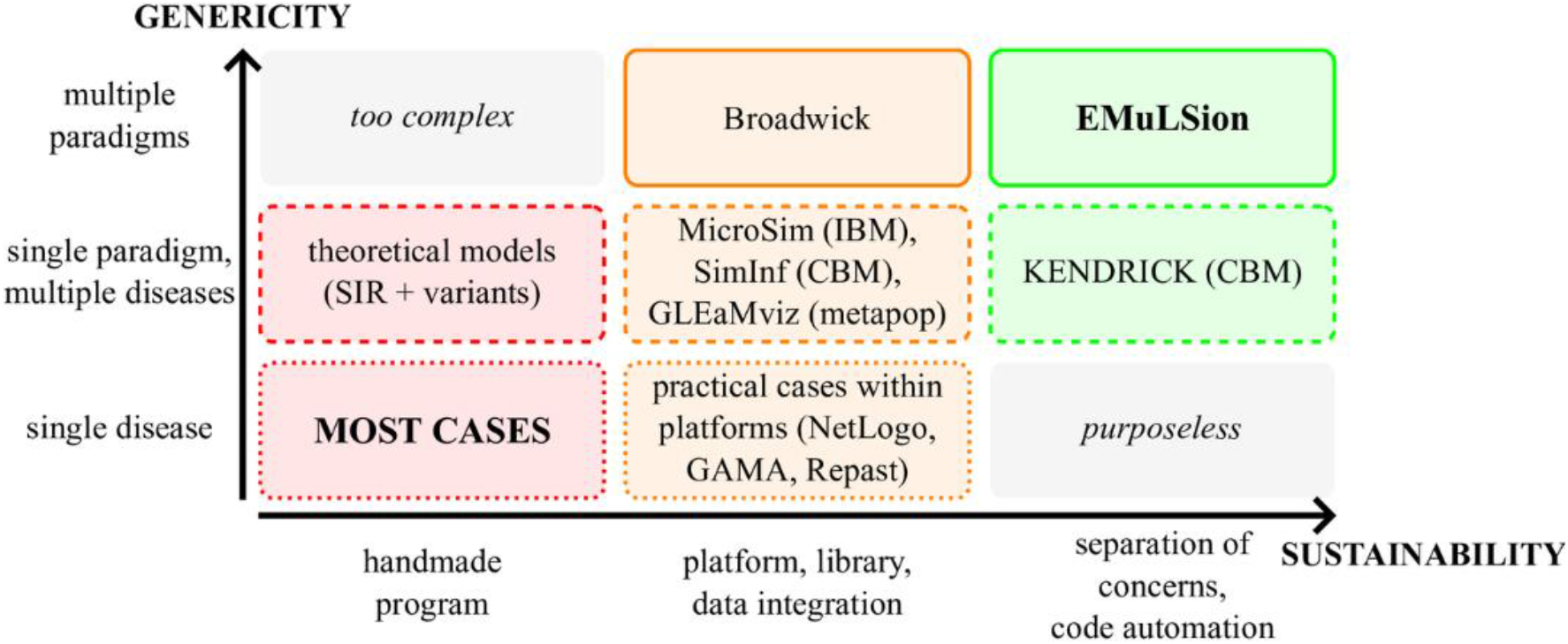
A taxonomy of modelling approaches in computational epidemiology. The vertical axis is based on the scope (from a single disease to multiple modelling paradigms); the horizontal axis represents the level of computer science complexity involved (from ad-hoc monolithic programs to a full separation between explicit knowledge and simulation code). In EMULSION, the coupling of a modular simulation architecture with a DSL is beneficial on both levels.

## Methods

### Coupling a Domain-Specific Language with a generic simulation engine

Designing a model using EMULSION involves three interdependent elements: 1) an explicit, modular and readable description of the model written using EMULSION’s DSL — this step is intended to be accessible to non-computer scientists experts; 2) the use of the generic simulation engine written by computer scientists, to capitalize, in an extensible, modular and reliable way, recurrent treatments and calculations that can be found in most epidemiological models (e.g. computation of states evolution over time, connection to data, etc.) — this engine is aimed at building and running the appropriate simulation architecture based on the model specifications in the DSL; 3) small code add-ons which may be necessary to add features (calculations, actions, processes) either specific to each model or not yet incorporated into the generic engine (Fig 2). The complexity of designing and implementing a model is thus broken down into several simpler concerns, without unnecessary code writing. Besides, models described through EMULSION’s DSL are univocal in the sense that they have to make most assumptions explicit, and refer to the simulation methods implemented within the generic engine.

**Fig 2.**
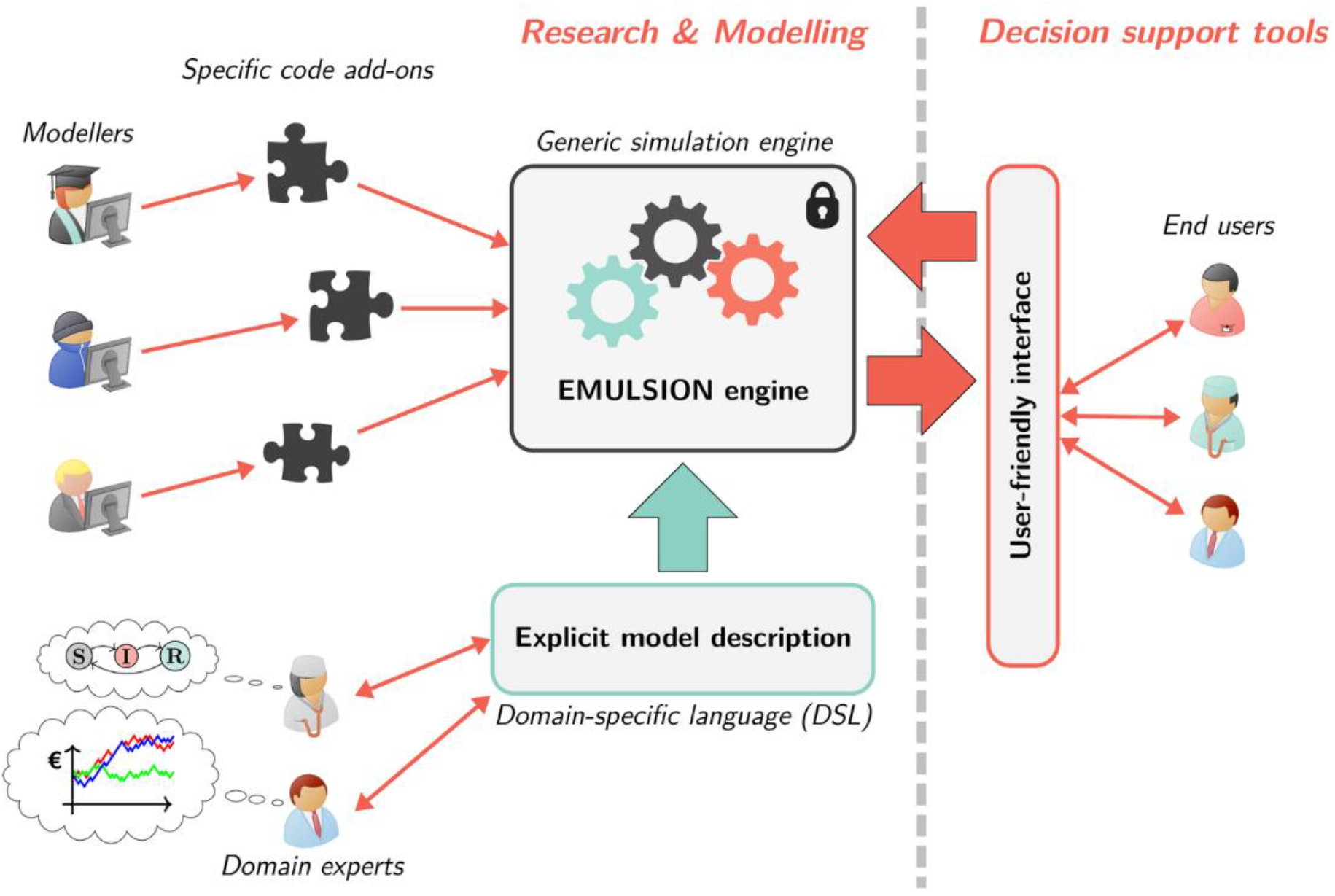
Approach enforced in EMULSION. A generic simulation engine is coupled to a domain-specific modelling language, fostering continuous experts’ involvement and user-friendly interactions. Experts’ knowledge is kept explicit, understandable, and revisable. A few specific code add-ons can be written to complement the simulation engine.

### Knowledge engineering: a paradigm-independent representation of processes

Epidemiological models mainly rely on the description of infectious processes. As a balanced formalism, we propose to extend flow diagrams to represent state evolution through Finite State Machines [43], widespread used in computer science. Features that were implicit in epidemiological design can be described explicitly in nodes (states) and edges (transitions) of state machine diagrams (Fig 3), enhancing the intelligibility of models. States can be endowed with 1) a duration distribution, specifying how long an individual is expected to stay in the current state, and 2) actions performed by individuals when entering, being in, or leaving the state. Transitions are labeled with either a rate, a probability or an absolute amount; they can also specify: 1) calendar conditions to indicate time periods when transitions are available; 2) escape conditions allowing to free from state duration constraints; 3) individual conditions to filter who is allowed; 4) actions performed by individuals crossing the transition (after leaving their current state and before entering their new one).

**Fig 3.**
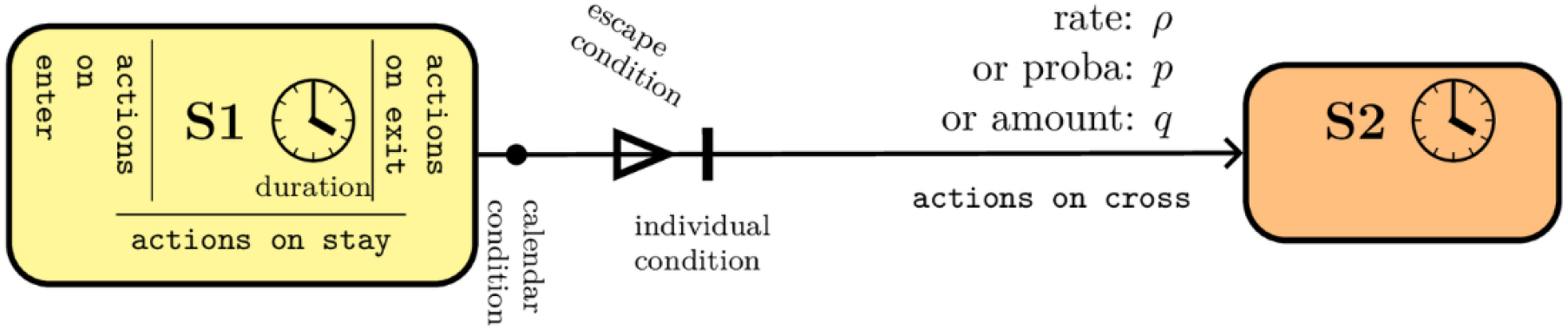
Structure of a transition between two states in state machines. States can be given a duration and actions when entering, staying in, or leaving the state. Transitions feature a rate, or probability, or amount, and can be associated with actions performed on crossing, time-dependent (“calendar”) conditions, or individual conditions restricting the capability to cross the transition, and escape conditions allowing individuals to leave their state before the nominal duration.

A classical flow diagram is essentially a conceptual sketch of the model, requiring further programming to control transitions between states; conversely, the state machine diagram is informative enough to be processed directly by the generic simulation engine without further code writing. For instance, flow diagrams generally assume an exponential distribution of durations in health states; but, sometimes other distributions are required, e.g. a constant incubation duration. While a classical approach would require to rewrite the simulation code to switch from exponential to constant duration, with EMULSION the specification of a constant duration in the node of the state machine is automatically handled with the correct computation by the generic engine. Similar changes can be made or revised on the fly, since they require no more than adding or removing a few lines in the model description.

Each state machine is aimed at describing a single process (infection, demography, …). Thus, instead of mixing concerns within a single, complex diagram, each process can be expressed, assessed, revised independently from the others and in a simple representation. Possible interactions between processes can be expressed through actions: for instance, actions performed in a ‘Treated’ state of a treatment process can induce changes in the infectious process.

While flow diagrams were population-oriented (i.e. describing the evolution of group size), state machines are individual-oriented: they specify individual behaviors, allowing to focus on fine-grained individual features. The subsequent issue, consisting in aggregating individuals to the relevant detail level without excessive computational cost, is addressed by the agent-based simulation architecture described a few lines below.

### A language for epidemiological model description

Model assessment, from the first assumptions to the interpretation of simulation results, is a long process. To keep the model explicit, understandable, and revisable throughout, it must be accessible under a readable form, rather than buried into the simulation code. Thus, we recommend gathering all model components (parameters, distributions, functions, time management, state machines, levels, processes occurring on each scale, actions, etc.) within a structured text file. We defined a Domain-Specific Language (DSL) [44] matching the needs of epidemiology, to allow experts to structure models through key-value pairs. Model description files are intended to be comprehensive documents, thus force modelers to leave comments and sources for each item: the same file can then be used to produce figures, parameter tables, or technical documentation. When processing model files, the generic engine parses parameters, variables, and mathematical expressions using a symbolic computing library, to translate them into true functions. It also builds the simulation architecture and checks the consistency of the model before running the simulation.

This separation between experts’ and domain-specific knowledge (declarative part of the model) on the one hand, and the algorithms to handle it (procedural part) on the other hand, is a classical, but powerful Artificial Intelligence solution [45], known for helping experts to be involved directly in the model design process, and for allowing fast, iterated feedbacks. Besides, this approach appears a kind of “literate modelling” by analogy with Knuth’s “literate programming” approach [46], aimed at fostering a human-friendly, purpose-driven way of developing software codes. The elaboration of a DSL for epidemiological models is actually a first attempt towards standardization, which must be supported by an ability to encompass existing modelling paradigms and adapt to real use cases.

The modelling language defined in EMULSION is an “internal” DSL [47], as it is based on another language, YAML (a human-friendly data serialization standard). Its syntax is quite simple, relying mainly on lists and on dictionaries (i.e. key-value pairs), which can be nested one in another and, for most components, do not require a special ordering. Contrary to most general-purpose programming languages, this DSL is aimed at describing declarative knowledge, i.e. the model components and their relations, the way to process them being implemented in the generic simulation engine. A whole example is provided as supporting information (with syntactic colorization: Additional File, S4 Files). Six main sections (first-level keys in the dictionary) must be specified: 1) the levels of interest in the simulation (e.g. individuals, populations, metapopulations…) and their link to agent classes (i.e. either agents defined in the generic simulation engine, as described below, or derived from the latter to provide specific code add-ons); 2) the processes occurring at each level, which can be either handled through a state machine (e.g. infection process, population dynamics, etc.), or implemented as a specific code add-on in the class associated to the level; 3) the description of the state machines, composed of the list of the states, with associated duration and actions if any, and the list of transitions between states, with possible conditions and actions (the description of the state machines is equivalent to the state machine diagram presented on Fig 3); 4) the comprehensive list of parameters used in the model, with their description and values; 5) the list of agent variables (“statevars”) with their role; 6) the list of agent actions with their description. Only the items of the two latter require subsequent implementation in the proper agent classes as specific code add-ons. Additional features can be specified in the model, such as time management (e.g. time unit, duration of discrete time steps, scheduling of events…) or desired outputs.

The key point is that this description, which can be developed and consulted independently of the generic simulation engine and of any possible code add-ons, does not require any computer science skill to be understood and discussed. Hence, it fosters interactions with experts throughout modelling, from formulating initial assumptions to specifying parameter values and identifying relevant scenarios and outputs. Besides, an EMULSION model not only enforces an explicit specification of model items that otherwise would be hidden in the code, but also requires a textual description of their rationale and purpose, to keep track for instance of the evolution of assumptions or of the exact meaning of parameters. Revising the model to account for new knowledge or to test alternative hypotheses essentially consists in modifying the YAML file, by adding or removing states, transitions, parameters, processes or actions, as shown on the application to Q Fever disease in the Results section.

### An agent-based software implementation

Several elements in the model description file rely upon the agent-based software architecture used in EMULSION, which is instantiated at runtime by the generic simulation to build the actual simulation from the model description.

Multi-Agent Systems (MAS) [48, 49] have become a classical paradigm for the simulation of complex systems. Agents, endowed with behaviors reflecting assumptions of a mechanistic model, interact within a shared environment. They are quite flexible and can represent any kind of entity, since they are defined by their behaviors and interaction capabilities rather than by their structure. Their behaviors can be defined through rules, equations, probabilistic trials, etc. More recently, in multi-level MAS, agents are also used to explicitly represent intermediary organization levels within the system, with their own behavioral capabilities. Hence, they can be used to encapsulate other paradigms within a homogeneous interface. Among the few generic meta-models designed for multi-level agent-based simulation [50–53], we used the principles defined in [50] which proved flexible enough to adapt to other fields [54], and provide useful features for computational epidemiology, such as a clear separation between declarative and procedural aspects. A multi-level MAS, in this meta-model, is the combination of an architecture of nested agents (through a hosting relation) and an explicit and intelligible description of the processes where agents are involved.

State machines specify quite accurately individual behaviors for each possible process. In most situations however, keeping all individuals in the simulation would be highly inefficient and lack relevance. Therefore, agents can materialize groups at different levels, according to model requirements and to similarities between individuals. EMULSION intends to implement those principles, by combining agent classes defined to match typical relationships between a group and the underlying entities [55]. All agents are situated in at least one environment where they live, perceive, and act. Two main agent families are used: atoms representing individuals, and groups representing aggregation with a tunable granularity level.

Depending on how such agents are combined and whether or not their behavior is controlled by a state machine, classical epidemiological modelling paradigms can be easily reproduced. A “GroupManager” agent endowed with a health-related state machine and hosting several “Compartment” agents reproduces a CBM (Fig 4, right). Contrarily, gathering “EvolvingAtom” agents, each one owning a state machine, within a “SimpleView”, leads to an IBM (Fig 4, left). Refer to Additional File, §S1 Appendix and S1 Fig for the detailed relationships between agent classes.

**Fig 4.**
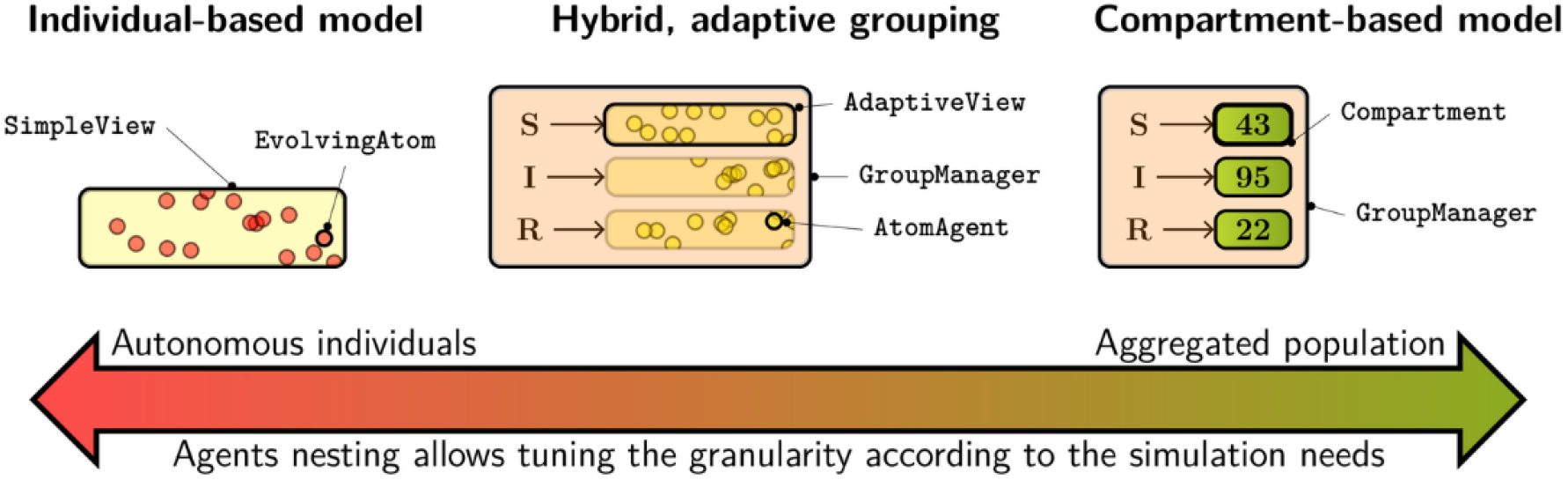
Integration of multiple modelling paradigms and scales. EMULSION allows multiple modelling paradigms to be expressed within the same formalism, based on nested agents, from the explicit representations of individuals (individual-based models) to aggregated populations (compartment-based models), with intermediary representations designed to group individuals depending on domain-dependent variables.

In addition, multi-level agents enable a hybrid modelling paradigm, mixing the preservation of individual states as in IBM and the reduced computational cost of CBM. Indeed, aggregation structures can be built at an intermediary stage, between individual- and population-oriented architectures. Individuals can indeed be gathered according to each separate concern, based on their similarities regarding each key variable. For instance, since the description of the infectious process is associated with a specific state machine, each atom agent can be endowed with a variable for holding this state (e.g. “health_state”). Then, it makes sense to gather individuals according to possible values of this “health_state” variable, for instance using “AdaptiveView” agents, hosted by a “GroupManager” (Fig 4, center). This “GroupManager” supervises the health-related state machine (instead of atoms) and, during the simulation, determines how many atoms have to change their “health_state” value and move from one “AdaptiveView” to another. To do so, due to the homogeneity of atoms within each “AdaptiveView” regarding “health_state”, only one multinomial sample per group is required, instead of one Bernoulli trial per individual, which reduces significantly the computation cost compared to a classical IBM. Besides, using “AdaptiveView” agents as containers facilitates the separation of concerns: if another process (e.g. recovery) suddenly affects the “health_state” variable of some agents, their change is detected by the view, which asks its own host (the “GroupManager”) to move modified atoms to the proper place.

Metapopulation appears a gathering of lower-level agents, such as those built after one of the previous architectures. “MultiProcessManager” agents are designed to host them, provide a contact structure, and be automatically constructed by EMULSION with the underlying components, based on the specification of processes modelling the contact structure, key variables and state machines in the model description file. Thus, several concerns are handled at the same time, without any special development effort for the model designer.

EMULSION models can be checked prior to simulation to identify missing or inconsistent information, and code templates can be generated to facilitate writing the specific add-ons. To run a simulation from a model, EMULSION parses the YAML description file to read parameters, resolve expressions, build the state machines, and instantiate the agent classes corresponding to the required levels and groupings, based both on the objects of the generic simulation engine and on the specific code add-ons.

### Application to the exploration of an epidemiological model (Q fever spread)

The algorithms within EMULSION have been broadly tested on several very well-known variants of SIR-like models [5, 6] based on CBM, IBM, hybrid modelling, and metapopulations. However, the major added-value of EMULSION is to facilitate the development of complicated models by model designers, to foster participative model revisions within short development time thanks to the DSL. Hence, we addressed models for a real disease, Q fever in dairy cattle herds, for which herd management processes have to be accounted for to reliably predict pathogen spread [15, 56, 57].

Q fever is a worldwide zoonosis caused by the bacterium *Coxiella burnetiid*. It has recently spread in Europe, e.g. in the Netherlands with a large number of human cases reported in 2007–2009 [58]. Domestic ruminants are recognized as the main source of human infection. In previous studies, a detailed individual-based within-herd model was designed to help better controlling *C. burnetii* spread in cattle herds with a particular attention paid to the diversity of transmission pathways and levels of pathogen shedding by infected hosts [15]. The main parameters of a simplified variant of this model were estimated from epidemiological data [56]. A study in the French department of Finistère revealed seroprevalence levels for 2697 dairy herd by enzyme-linked immunosorbent assay (ELISA) in bulk tank milk in 2012. 797 herds were detected seronegative in 2012 and tested again one year later. The annual herd incidence (number of herds newly infected) was of 295 herds. The within-herd model was extended to the between-herd level and confronted to such epidemiological data [57]. However, three main computational and epidemiological issues remained. First, the infection process was mixed with the reproduction cycle of cows, impeding modifications of biological assumptions and, thus, the exploration of a larger variety of model structures. Second, the integration of within-herd dynamics into the between-herd scale was not straightforward. Third, the simulated annual herd incidence was still too low.

We re-implemented these models using EMULSION to explore more quickly the interplay between within-herd and between-herd levels. First, the original model [15] was simplified, keeping relevant assumptions and removing those less crucial in the perspective of the between-herd dynamics [59]. Then, it was extended to the regional between-herd level, where assumptions regarding airborne transmission were compared. Finally, the within-herd model was revised with alternative hypotheses in the infection process, to better reproduce the observed annual herd incidence at the between-herd level while making plausible assumptions about host infection processes. The relative roles of trade and airborne transmission in regional pathogen spread were reassessed under these new modelling assumptions.

## Results

### Model simplification within EMULSION

According to the parsimony principle, we built a model with a minimum number of states, transitions and parameters, trying to reproduce main simulation outcomes (prevalence, seroprevalence and bacterial shedding) of the original within-herd model [15] after the transient period. To do so, we identified possible simplifications (reduction of the number of states and transitions, and replacement of distributions by aggregated parameters), and assessed them by simulation with a modified YAML configuration file, without changing the code, and iterated the process with alternative hypotheses. In the resulting model, called below “simplified model”, 5 (out of 11) health states were retained (Additional File, S2 Fig and S3 Fig): susceptible (S), infectious without (I−) or with (I+) antibodies, and carrier with (C+) or without (C−) antibodies, without distinguishing between shedding levels or pathways. Contaminations occurred through contacts with contaminated environment, bacteria (*E_totai_*, bacterial load in environment) coming either from local shedding (*E_local_*, due to infectious animals from the herd) or external sources (*E_aero_*, from airborne transmission). At herd scale, when neglecting between-herd transmission, *E_local_* is equal to *E_total_* (Additional File, S2 Appendix equation 1). The probability of infection was determined by 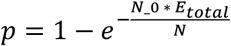, where N denoted the population present in the herd (N_0 being a normalization factor). Besides, only adult female cows were taken into account. They follow a reproduction process (Additional File, S4 Fig) with state depending on their pregnancy status (P for pregnant, NP for non-pregnant). Transitions between P and NP were handled by a duration in each state. Special events such as abortion could happen to pregnant cows up to three weeks after infection. Local bacteria shedding occurred either during infectious states through on-stay actions, or massively at special events (calving and abortion), through on-cross actions. The YAML file corresponding to this model (model structure, levels, parameters, variables, etc.) is provided as supporting information (Additional File, S4 Files).

The model followed the hybrid structure (Fig 4, center) to fully account for individual events (calving and abortion) without excessive computational cost. The implementation required only a class for individuals, derived from “AtomAgent”, and one for the herd, derived from “MultiProcessManager” [59]. The resulting multi-agent architecture used to represent a herd is shown on Fig 5. Processes involved at the herd level were the following: 1) culling (cow removal depending on parity); 2) replacement (introduction of new animals); 3) infection and 4) farm management, both based on state machines respectively affecting health state and life cycle; 5) actualization of animal grouping by parity; 6) bacterial decay in the environment (exponential decrease) and update of bacterial shedding. While processes 1, 2, and 6 required writing short specific code add-ons, the others, involving generic mechanisms such as state machines and groupings based on parity, health state, and life cycle, were handled automatically by the generic simulation engine. Parameters were calibrated to match the median and 10-90 percentiles (on 200 repetitions) of three major outcomes (prevalence, seroprevalence and bacterial load in environment) of the original model simulation after the transient period (200 weeks after introducing one I+ cow just before calving in a fully susceptible herd).

**Fig 5.**
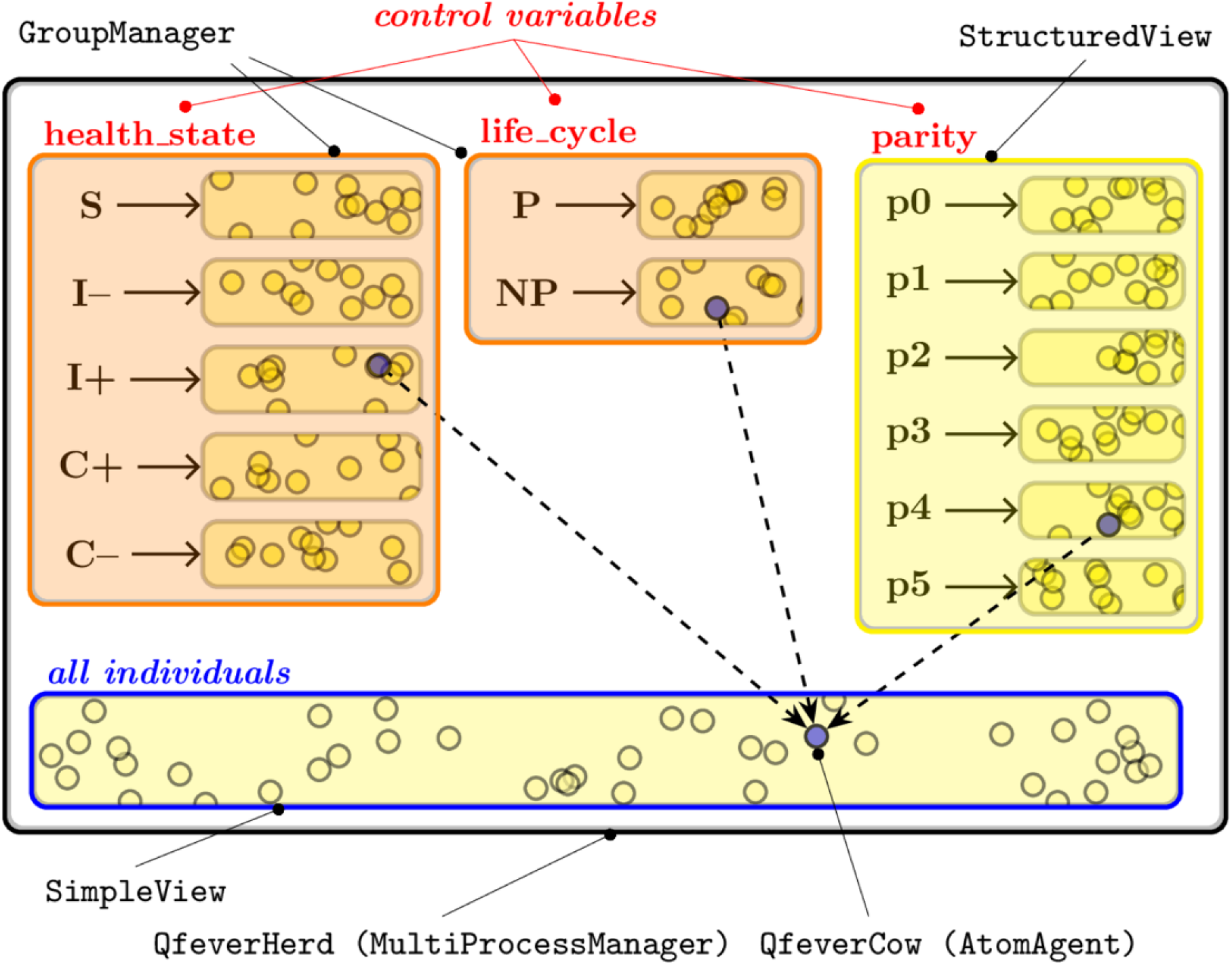
Structure of the within-herd hybrid model for Q fever. Individuals are aggregated according to concern-related variables (health state, life cycle, and parity). Individuals (e.g. the blue one in health state I+, life state NP, and parity 4) can be accessed through each concern or through a global list.

### Exploration: from within-herd to between-herd levels, back and forth

Next, we focused on accounting for annual herd incidence observed in Finistère. The between-herd model is a metapopulation, composed of independent herds linked through a contact network. As in the within-herd architecture, the metapopulation could be implemented in EMULSION by a “MultiProcessManager” agent, encapsulating a view holding all herds and endowed with dedicated processes to handle interactions between them (Additional File, S7 Fig). Transforming the YAML file from the within-herd to the between-herd scale only required to add the description architecture and processes at the metapopulation level (Additional File, S1 Text), here assuming herds have similar parameter values and sizes.

Herds could interact either through animal trade or airborne transmission from neighbor herds. Initial assumptions considered animals bought from outside the metapopulation healthy. Bacteria were transported and deposited by wind using a plume dispersion equation [60, 61] (called “Ermak-Stockie function” below and detailed in Additional File, §S2 Appendix). Processes involved at the metapopulation level thus were: 1) activation of herd processes; 2) airborne transmission; 3) selection of animals for trade movements in source herds; 4) effective movement to destination herds. Herd specificities (initial size, renewal, culling and trade movements) were based on the French livestock exchange data, requiring specific code add-ons to make the metapopulation agent calibrate herd parameters and extract the relevant trade movements from data. The predicted herd incidence with these initial assumptions was much lower than in observed data (Fig 6-A). The discrepancy between observed and predicted incidence could be explained either by a missing transmission route (but no other is known for Q fever), wrong assumptions about risky trade or airborne transmission, or wrong assumptions about the within-herd infection dynamics. To check for these two latter issues and improve herd incidence predictions, we considered alternative assumptions on three main levers. We first assumed that animals coming from outside the metapopulation had the same probability of being infected as inside the metapopulation (rather than assuming them susceptible), this one being a part of a larger regional population of herds. The impact on herd incidence was low (Fig 6-B) despite 18% of incoming movements from outside the metapopulation. Second, regarding airborne transmission, we assumed that individuals could be contaminated by inhalation of available bacteria (which is biologically plausible) rather than by deposited pathogens. Hence, a simpler Gaussian approach [61], not accounting for deposition, was implemented for plume dispersion (Additional File, equation 6, §S2 Appendix). Only a few lines of codes were changed in the specific add-on to define the new function. Still, parameters could not be calibrated within biologically plausible ranges to reach the expected seropositive herd incidence (Fig 6-C), while the true herd incidence sharply increased. The difference between incidence levels considering either infectious or seropositive animals, suggested to investigate the within-herd model further. We reexamined shedding assumptions, shedding being observed to be intermittent [62]. The original model [15] assumed that I− cows were able to eliminate all bacteria and become S again (non-shedder without antibodies and then apparently susceptible) [56], resulting in transitions from I-to S. Alternatively, a latent state (L) could have been assumed, i.e. a non-shedding but infected state. The intermittent shedding then can be explained by a loop between L and I-states (Additional File, S5 Fig), obviously increasing within-herd prevalence and reducing sharply spontaneous fade-out at local scale. Using EMULSION, going back and forth from one model structure to another is straightforward, even for a multi-scale model. The new within-herd model (“Latent state model”) was built from the Simplified model by adding a state and changing four transitions in the state machine describing health states (i.e. 10 lines in the YAML file: Additional File, S2 Text). It was then calibrated to keep the same steady-state regarding the same three main simulation outcomes (prevalence, seroprevalence and bacterial load in environment) in the medium run as the simplified model (Additional File, S6 Fig). Then, back to between-herd scale, we calibrated parameters in a plausible range for reaching the expected herd incidence level (Fig 6-D).

**Fig 6.**
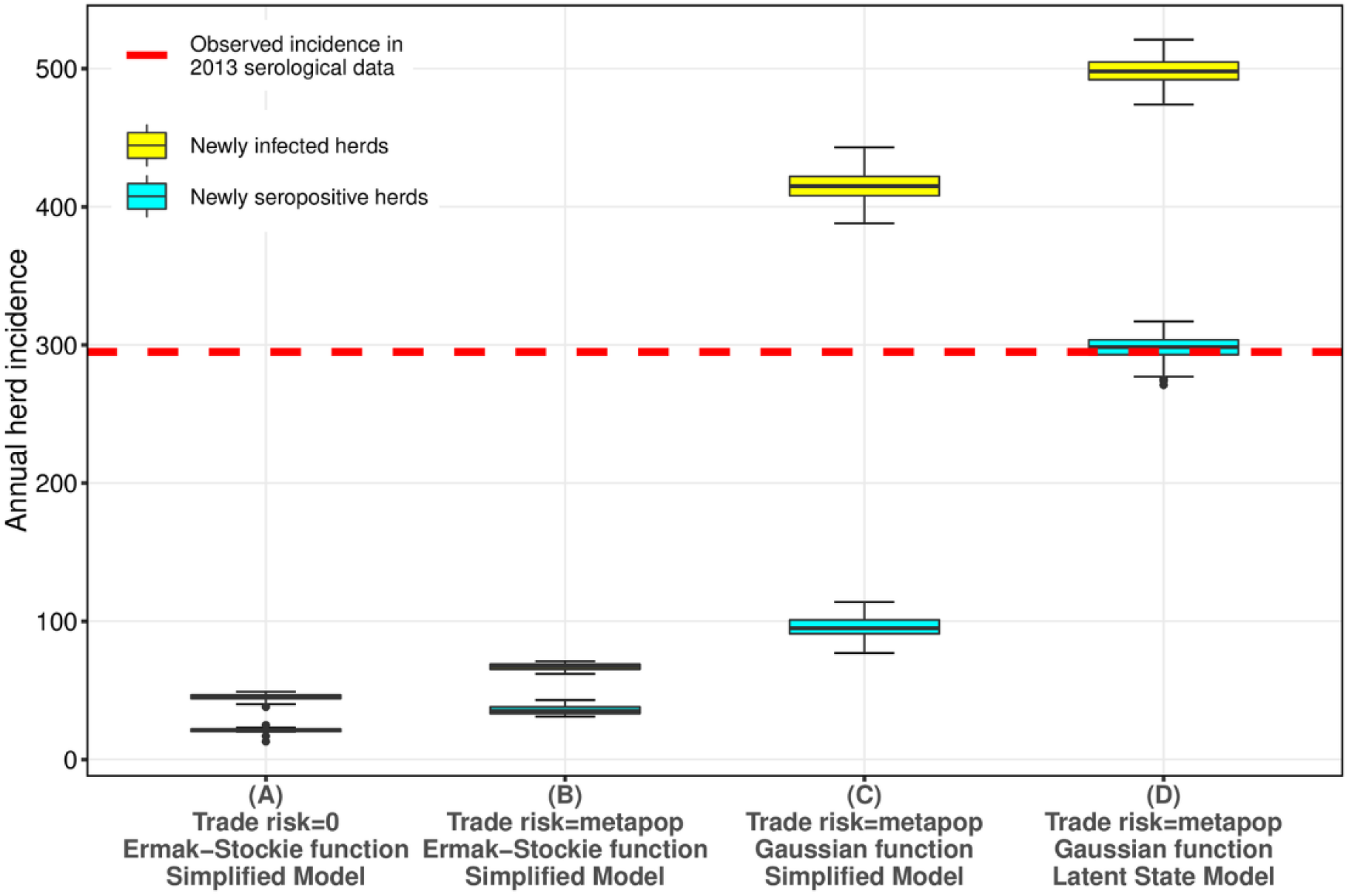
Annual herd incidence distributions in models based on combining variants of three main features. 1) No infection risk when purchasing animals from outside the metapopulation vs. risk similar to the average prevalence in the metapopulation, 2) Airborne transmission calculated by the Ermak-Stockie function vs. a Gaussian function, 3) Simplified within-herd model vs. model with a latent state. Yellow (cyan) color corresponds to the number of herds observed healthy (seronegative) in 2012 that hold a shedder (seropositive) animal at least once during the year. Observed data correspond to the 295 herds newly detected as seropositive by the ELISA test in 2013 (seronegative in 2012) in Finistère, France.

A sensitivity analysis was carried out to assess the impact of model parameters on two main outcomes at the metapopulation level (annual and weekly herd incidence), highlighting *N*_0_ (normalization factor), *l* (transition probability from L to I− states), *m* (transition probability from I− to S), κ (proportion of bacteria leaving local environments to contribute to airborne transmission) and ζ (contact rate with bacteria coming from airborne transmission) as key parameters (Additional File, S8 Fig). In addition, we ensured that spatial distributions of herds by health status were similar between observed serological data and simulations (Additional File, S9 Fig).

### Impact on previous conclusions

Regarding pathways responsible for Q fever spread at the regional scale, we knew from the previous study [57] that airborne transmission was predominant over trade movements. Yet, we wanted to assess whether trade-borne infections could be neglected or not. After exploring model assumptions and parameters to account for observed seroprevalence data in 2013 (Fig 6), we examined more finely temporal and spatial effects of both transmission pathways, under the hypothesis that seroprevalence is a relevant indicator for disease persistence within a one-year interval. First, infections of naive herds appeared, as expected from previous results, to be caused mostly (89.5% of newly infected herds) by airborne transmission (Fig 7, A). However, it also appeared that contaminations caused by trade movements happened in a more deterministic way, which was not highlighted previously. When we considered the infection dates of herds contaminated by trade movements at least once in 50 stochastic repetitions by trade movements, two groups were identified (Fig 7, B): first, herds infected early in the simulations, almost always at the same date and by trade movements; second, herds infected at a variable date and with a variable contribution of airborne transmission. Spatial distribution of incident herds (Fig 7, C) pointed out that areas with a high density of initially infected herds (and of herds in general: Additional File, S10 Fig) drove at the same time the predominance of airborne transmission and the probability that a herd subject to airborne transmission risk becomes infected. Conversely, herds infected at least once by trade movements were mostly located on peripheral areas, such as coasts (Fig 7, D), and most of them purchased animals directly in an initially prevalent herd before becoming infected, with mostly early infection dates. To summarize, the spread of Q fever within areas of high prevalence and high herd density is strong and mainly caused by airborne transmission, which argues for vaccination as a disease control strategy in such areas, while herds in low prevalence areas have little chance to be contaminated but by trade, which supports tests on purchase in that case.

**Fig 7.**
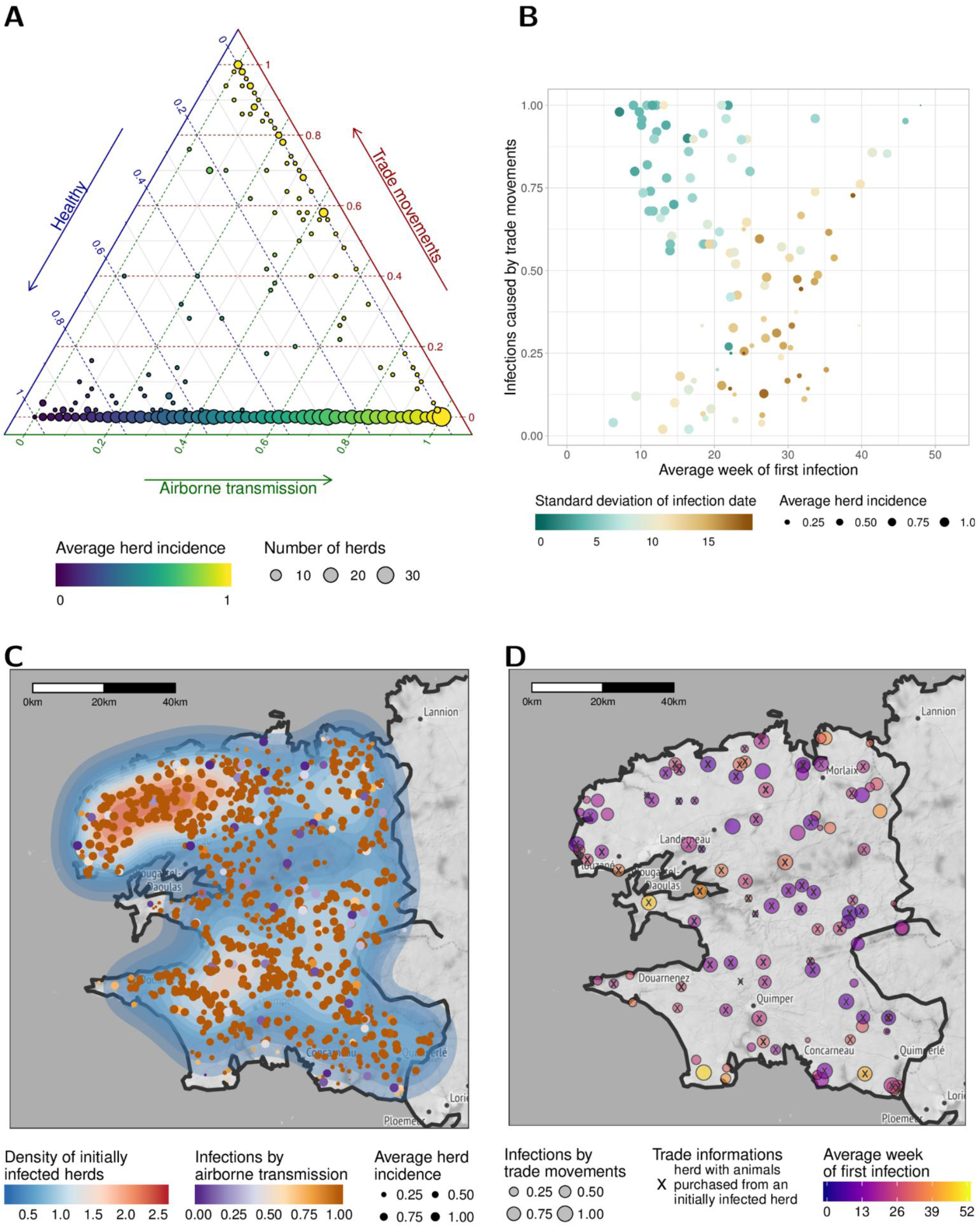
Contributions of infection pathways at regional scale, as predicted by the between-herd Q fever model with latent state and a Gaussian plume airborne transmission (50 stochastic repetitions of the standard scenario). (A) Distribution of the proportion of repetitions where each herd stayed healthy, became infected by animal trade movements, or became infected by airborne transmission, over one year (color shows the proportion of repetitions where the herd became infected by any of the transmission routes). (B) Amongst herds infected in at least one repetition by trade movements, relation between the proportion of infections caused by movement and the infection date (in average and standard deviation), exhibiting two subgroups: one with an early and little variable infection date, caused very often by movements (blue points), and the other with a more variable infection date and caused less often by movements (brown points). (C) Map of Finistère with the density of initially infected herds (2012 serological data) and the location of herds infected in at least one repetition. Color shows the proportion of infections caused by airborne transmission vs. trade movements. (D) Map showing the location and average infection date of herds infected at least once by trade movements. Herds marked with a “x” purchased animals from initially infected herds before their own infection. Map tiles by Stamen Design, under CC BY 3.0. Data by OpenStreetMap, under ODbL.

## Discussion and conclusions

EMULSION is the first framework that simultaneously provides a Domain-Specific Language dedicated to the comprehensive and accurate description of epidemiological models, from SIR-like models to more complex multi-scale multi-concern models, together with a modular simulation engine using a multi-level agent-based architecture, to encompass existing epidemiological issues and modelling paradigms within a homogeneous interface (Fig 1).

Elaborating realistic models (such as [15, 57]) often requires many trials aiming at the exploration of various assumptions and processes. EMULSION significantly accelerates model development, first because it provides classical computational bricks, but also since changing hypotheses (e.g. adding or deleting a state or a transition) generally consists in modifying the configuration file, instead of rewriting many parts of a large specific source code. The modularity of model description allows assessing separately hypotheses, which demands much more work when dealing with ad-hoc models and is more prone to programming and hence interpretation errors.

Using EMULSION, we re-implemented very quickly the initial within-herd Q fever dairy cattle model and assessed the compatibility of simpler alternative model structures with previous predictions derived from a model whose parameters were estimated using observed on-farm data. Then, moving to the regional scale required little transformation of the within-herd model. Small pieces of code related to trade and weather data and calculation of transmission functions were used as specific add-ons by the simulation engine. The facility to compare a large panel of assumptions regarding the existence of a latent stage in Q fever infectious process, the nature of airborne transmission, and values of the less well-known parameters, allowed us to identify and calibrate the best candidates with respect to observed herd incidence data. Also, conditions under which trade movements and airborne transmission contribute to new infections were explored, highlighting new findings, especially that infections caused by movements are almost deterministic and impact mostly herds in peripheral areas and low prevalence areas.

This works provides a proof of concept, demonstrating the added-value of using such a framework, both in terms of code reduction and model readability. To promote our approach and modelling language, EMULSION will be soon released as an open-source software. The current version of the generic simulation engine being written in Python, efficiency cannot compete with compiled languages such as C++ (as the code generated by KENDRICK [40]) or Java (used in the Broadwick framework [38]), especially at the between-herd scale. This was not under the scope of the present study. Nevertheless, the facility to choose the granularity level of simulations and the adaptive gathering of individuals make the approach much more efficient than IBM anyway. To tackle efficiency issue, the next step will be to consider using EMULSION’s DSL to build dedicated, optimized code from model descriptions.

We showed how modelling paradigms and scales could be wrapped within agents, which are in charge of processing the required calculations according to their specificities. This allows modellers to focus on their research questions instead of implementation issues, while still being able to select and compare relevant modeling paradigms. To go further, new computational issues in epidemiological modelling have to be addressed, especially in coupling contrasted paradigms [63]. For instance, multihost pathosystems may combine populations having highly contrasted characteristics, such as size (a large size leading to more deterministic dynamics while a small one enhances stochastic events) and movement patterns (vectors spread continuously while livestock trade and human activity give rise to discrete long-distance jumps). In addition, the level of required details may differ, possibly leading to combine aggregated representation (i.e. CBM) with preservation of individuals (i.e. IBM). The integration of such features into both the DSL and the generic engine (especially as new agent classes) will enable modellers to address such computational challenges in epidemiological modelling.

From the point of view of computer science, adapting the original multi-level agent-based metamodel [41, 50] to epidemiological issues also was fruitful. Dealing with aggregation and disaggregation in an adaptive way is a challenging open question in multi-level modelling. The architecture designed for multi-scale epidemiological systems will provide clues for building similar structures in general-purpose agent-based systems, and continue the identification and characterization of design patterns in multi-level agent-based simulation initiated in [55].

We consider our contribution a first step towards a standardized DSL for epidemiology. Though initiated in the context of animal health, our approach is generic and modular enough to extend to human and plant epidemiology. The generalization of such methods could enhance significantly the reactiveness of modelers in sketching, assessing, and recommending reliable and efficient control measures against outbreaks, accounting for possible biases in model predictions arising from uncertainty in model assumptions.

## Supporting information

## Additional material

**Additional File. (PDF) Complementary information for the main article**. Contains a description of agent classes, of airborne transmission functions, additional figures, and YAML file descriptions or modifications.

## Abbreviations

CBM: Compartment-Based Model;
DSL: Domain-Specific Language;
IBM: Individual-Based Model;
MAS: Multi-Agent Systems;
ODD: “Overview, Design concepts, Details” protocol;
ODE: Ordinary Differential Equations;
SBGN: Systems Biology Graphical Notation

## Declarations

### Ethics approval and consent to participate

Not applicable.

### Consent for publication

Not applicable.

### Availability of data and material. Data

French livestock exchange data were provided by the French Ministry of Agriculture (FMA). Data collection and analyses are subject to a confidentiality agreement (available upon request from the following contact point). Public weather datasets were provided by the European Centre for Medium-Range Weather Forecasts (ECMWF). Parameters and structure of Q Fever models are fully available in Supporting Information.

### Software

The framework EMULSION is awaiting approval from the French National Institute for Agricultural Research (INRA) before being publicly released. In the meanwhile, the specific code add-ons used for Q fever modelling and the generic simulation engine are available upon request from the contact author.

### Authors’ contributions

SP designed the DSL and the engine of EMULSION, led software developments, and drafted the manuscript; YLH participated in software development, carried out Q fever study, and drafted the corresponding section; VS participated in the development and data preparation; TH, EV and FB provided expertise on Q fever and participated in results interpretation; PE supervised the specifications of EMULSION, and designed the Q fever study. All authors helped draft the manuscript and gave final approval for publication.

## Acknowledgments

We are grateful to the French Ministry of Agriculture for granting us access to cattle datasets.

## References

1. Peng RD. Reproducible Epidemiologic Research. Am J Epidemiol. 2006;163:783–789.

2. Sandve GK, Nekrutenko A, Taylor J, Hovig E. Ten Simple Rules for Reproducible Computational Research. PLoS Comput Biol. 2013;9:e1003285.

3. Leek JT, Peng RD. Opinion: Reproducible research can still be wrong: Adopting a prevention approach. Proc Natl Acad Sci. 2015;112:1645–1646.

4. Kermack WO, McKendrick AG. A Contribution to the Mathematical Theory of Epidemics. Proc R Soc. 1927;:700–721.

5. Diekmann O, Heesterbeek HJ. Mathematical epidemiology of infectious diseases: model buildign, analysis and interpretation. Chichester: Wiley; 2000.

6. Keeling MJ, Rohani P. Modeling Infectious Diseases in Humans and Animals. Princeton University Press; 2008.

7. Merali Z. Computational science: … Error. Nature. 2010;467:775–777.

8. Hethcote HW. The Mathematics of Infectious Diseases. SIAM Rev. 2000;42:599–653.

9. Marcé C, Ezanno P, Seegers H, Pfeiffer D, Fourichon C. Predicting fadeout versus persistence of paratuberculosis in a dairy cattle herd for management and control purposes: a modelling study. Vet Res. 2011;42:36.

10. DeAngelis DL, Grimm V. Individual-based models in ecology after four decades. F1000Prime Rep. 2014;6. doi:10.12703/P6-39.

11. Railsback SF, Grimm V. Agent-Based and Individual-Based Modelling: A Practical Introduction. Princeton University Press; 2011.

12. Ferguson NM, Cummings DAT, Cauchemez S, Fraser C, Riley S, Meeyai A, et al. Strategies for containing an emerging influenza pandemic in Southeast Asia. Nature. 2005;437:209–14.

13. Halloran ME, Ferguson NM, Eubank S, Longini IM, Cummings DAT, Lewis B, et al. Modeling targeted layered containment of an influenza pandemic in the United States. Proc Natl Acad Sci. 2008;105:4639–44.

14. Amouroux E, Desvaux S, Drogoul A. Towards Virtual Epidemiology: An Agent-Based Approach to the Modeling of H5N1 Propagation and Persistence in North-Vietnam. In: Bui TD, Ho TV, Ha QT, editors. 11th Pacific Rim Int. Conf. on Multi-Agents (PRIMA). Springer; 2008. p. 26–33. doi:10.1007/978-3-540-89674-6_6.

15. Courcoul A, Monod H, Nielen M, Klinkenberg D, Hogerwerf L, Beaudeau F, et al. Modelling the effect of heterogeneity of shedding on the within herd *Coxiella burnetii* spread and identification of key parameters by sensitivity analysis. J Theor Biol. 2011;284:130–141.

16. Robins J, Bogen S, Francis A, Westhoek A, Kanarek A, Lenhart S, et al. Agent-based model for Johne’s disease dynamics in a dairy herd. Vet Res. 2015;46.doi:10.1186/s13567-015-0195-y.

17. Marshall BDL, Galea S. Formalizing the Role of Agent-Based Modeling in Causal Inference and Epidemiology. Am J Epidemiol. 2014;181:92–99.

18. Parker J, Epstein JM. A Distributed Platform for Global-Scale Agent-Based Models of Disease Transmission. ACM Trans Model Comput Simul. 2011;22:2:1–2:25.

19. Gilpin M, Hanski I, editors. Metapopulation Dynamics: Empirical and Theoretical Investigations. Elsevier BV; 1991. doi:10.1016/b978-0-12-284120-0.50003-6.

20. Grenfell B, Harwood J. (Meta)population dynamics of infectious diseases. Trends Ecol Evol. 1997;12:395–399.

21. Keeling M. The implications of network structure for epidemic dynamics. Theor Popul Biol. 2005;67:1–8.

22. Arino J, van den Driessche P. Disease spread in metapopulations. In: Brunner H, Zhao X-Q, Zou X, editors. Nonlinear Dynamics and Evolution Equations. American Mathematical Society; 2006. p. 1–12.

23. Beaunée G, Vergu E, Ezanno P. Modelling of paratuberculosis spread between dairy cattle farms at a regional scale. Vet Res. 2015;46.doi:10.1186/s13567-015-0247-3.

24. Ajelli M, Gonçalves B, Balcan D, Colizza V, Hu H, Ramasco JJ, et al. Comparing large-scale computational approaches to epidemic modeling: Agent-based versus structured metapopulation models. BMC Infect Dis. 2010;10.doi:10.1186/1471-2334-10-190.

25. Keeling MJ, Danon L, Vernon MC, House TA. Individual identity and movement networks for disease metapopulations. Proc Natl Acad Sci. 2010;107:8866–8870.

26. Gillespie DT. Exact stochastic simulation of coupled chemical reactions. J Phys Chem. 1977;81:2340–61.

27. Bretó C, He D, Ionides EL, King AA. Time series analysis via mechanistic models. Ann Appl Stat. 2009;3:319–48.

28. Le Novere N, Hucka M, Mi H, Moodie S, Schreiber F, Sorokin A, et al. The Systems Biology Graphical Notation. Nat Biotech. 2009;27:735–741.

29. Díaz-Zuccarini V, Pichardo-Almarza C. On the formalization of multi-scale and multi-science processes for integrative biology. Interface Focus. 2011;1:426–437.

30. Perra N, Balcan D, Gonçalves B, Vespignani A. Towards a Characterization of Behavior-Disease Models. PLoS ONE. 2011;6:e23084.

31. Grimm V, Berger U, Bastiansen F, Eliassen S, Ginot V, Giske J, et al. A standard protocol for describing individual-based and agent-based models. Ecol Model. 2006;198:115–126.

32. Amouroux E, Gaudou B, Desvaux S, Drogoul A. O.D.D.: A Promising but Incomplete Formalism for Individual-Based Model Specification. In: RIVF Int. Conf. on Computing and Communication Technologies. IEEE; 2010.doi:10.1109/rivf.2010.5633421.

33. Roche B, Guégan J-F, Bousquet F. Multi-agent systems in epidemiology: a first step for computational biology in the study of vector-borne disease transmission. BMC Bioinformatics. 2008;9.doi:10.1186/1471-2105-9-435.

34. Collier N, Ozik J, Macal CM. Large-Scale Agent-Based Modeling with Repast HPC: A Case Study in Parallelizing an Agent-Based Model. In: Parallel Processing Workshops (Euro-Par). Springer Nature; 2015. p. 454–465.doi:10.1007/978-3-319-27308-2_37.

35. Widgren S, Bauer P, Engblom S. SimInf: An R package for Data-driven Stochastic Disease Spread Simulations. ArXiv Prepr ArXiv160501421 Q-BioPE. 2016. http://arxiv.org/abs/1605.01421.

36. Cakici B, Boman M. A workflow for software development within computational epidemiology. J Comput Sci. 2011;2:216–222.

37. Broeck WV den, Gioannini C, Gonçalves B, Quaggiotto M, Colizza V, Vespignani A. The GLEaMviz computational tool, a publicly available software to explore realistic epidemic spreading scenarios at the global scale. BMC Infect Dis. 2011;11. doi:10.1186/1471-2334-11-37.

38. O’Hare A, Lycett SJ, Doherty T, Salvador LCM, Kao RR. Broadwick: a framework for computational epidemiology. BMC Bioinformatics. 2016;17. doi:10.1186/s12859-016-0903-2.

39. Haddad H, Moulin B, Thériault M. A fully GIS-integrated simulation approach for analyzing the spread of epidemics in urban areas. SIGSPATIAL Spec. 2016;8:34–41.

40. Bui T-M-A, Ziane M, Stinckwich S, Ho T-V, Roche B, Papoulias N. Separation of Concerns in Epidemiological Modelling. In: Fuentes L, Batory DS, Czarnecki K, editors. Proceedings of the 15th International Conference on Modularity. ACM; 2016. p. 196–200. doi:10.1145/2892664.2892699.

41. Picault S, Huang Y-L, Sicard V, Ezanno P. Enhancing Sustainability of Complex Epidemiological Models through a Generic Multilevel Agent-based Approach. In: Sierra C, editor. Proceedings of the 26th International Joint Conference on Artificial Intelligence (IJCAI’2017). Melbourne, Australia: AAAI; 2017.

42. Bobashev GV, Goedecke DM, Yu F, Epstein JM. A Hybrid Epidemic Model: Combining The Advantages Of Agent-Based And Equation-Based Approaches. Winter Simul Conf. 2007. doi:10.1109/wsc.2007.4419767.

43. Booth TL. Sequential Machines and Automata Theory. 1st edition. New York: John Wiley and Sons; 1967.

44. Mernik M, Heering J, Sloane AM. When and how to develop domain-specific languages. ACM Comput Surv. 2005;37:316–44.

45. Newell A, Shaw JC, Simon HA. Report on a General Problem-Solver Program. In: Proceedings of the International Conference on Information Processing. 1959. p. 256–264.

46. Knuth DE. Literate programming. Stanford, Calif.: Center for the Study of Language and Information; 1992.

47. Fowler M, Parsons R. Domain-specific languages. Upper Saddle River, NJ: Addison-Wesley; 2011.

48. Ferber J. Multi-agent systems: an introduction to distributed artificial intelligence. Harlow: Addison-Wesley; 1998.

49. Weiss G, editor. Multiagent systems: a modern approach to distributed artificial intelligence. Cambridge, Mass: MIT Press; 1999.

50. Picault S, Mathieu P. An Interaction-Oriented Model for Multi-Scale Simulation. In: Walsh T, editor. Proceedings of the 22nd International Joint Conference on Artificial Intelligence (IJCAI’2011). AAAI; 2011. p. 332–337. https://hal.archives-ouvertes.fr/hal-00826401.

51. Morvan G, Veremme A, Dupont D. IRM4MLS: The Influence Reaction Model for Multi-Level Simulation. In: Multi-Agent-Based Simulation XI. Springer; 2011. p. 16–27. doi:10.1007/978-3-642-18345-4_2.

52. Camus B, Bourjot C, Chevrier V. Multi-level modeling as a society of interacting models. In: Yilmaz L, Ören TI, Madey G, Sierhuis M, Zhang Y, editors. Agent-Directed Simulation Symposium (in SpringSim). SCS/ACM; 2013. http://dl.acm.org/citation.cfm?id=2499595.

53. Huraux T, Sabouret N, Haradji Y. A Multi-level Model for Multi-agent based Simulation: In: Proceedings of the 6th International Conference on Agents and Artificial Intelligence. SCITEPRESS - Science and and Technology Publications; 2014. p. 139–46. doi:10.5220/0004814501390146.

54. Maudet A, Touya G, Duchêne C, Picault S. DIOGEN, a multi-level oriented model for cartographic generalization. Int J Cartogr. 2017;3:121–33.

55. Mathieu P, Morvan G, Picault S. Multi-level agent-based simulations: Four design patterns. Simul Model Pract Theory. 2018;in press.

56. Courcoul A, Vergu E, Denis J-B, Beaudeau F. Spread of Q fever within dairy cattle herds: key parameters inferred using a Bayesian approach. Proc R Soc B Biol Sci. 2010;277:2857–65.

57. Pandit P, Hoch T, Ezanno P, Beaudeau F, Vergu E. Spread of *Coxiella burnetii* between dairy cattle herds in an enzootic region: modelling contributions of airborne transmission and trade. Vet Res. 2016;47. doi:10.1186/s13567-016-0330-4.

58. van der Hoek W, Morroy G, Renders NHM, Wever PC, Hermans MHA, Leenders ACAP, et al. Epidemic Q Fever in Humans in the Netherlands. In: Toman R, Heinzen RA, Samuel JE, Mege J-L, editors. Coxiella burnetii: Recent Advances and New Perspectives in Research of the Q Fever Bacterium. Dordrecht: Springer Netherlands; 2012. p. 329–64. doi:10.1007/978-94-007-4315-1_17.

59. Picault S, Huang Y-L, Sicard V, Beaudeau F, Ezanno P. A Multi-Level Multi-Agent Simulation Framework in Animal Epidemiology. In: Demazeau Y, Davidsson P, Vale Z, Bajo J, editors. Proceedings of the 15th International Conference on Practical Applications of Agents and Multi-Agent Systems (PAAMS’2017). Porto: Springer; 2017. p. 209–21.

60. Ermak DL. An analytical model for air pollutant transport and deposition from a point source. Atmospheric Environ 1967. 1977;11:231–7.

61. Stockie JM. The Mathematics of Atmospheric Dispersion Modeling. SIAM Rev. 2011; 53:349–72.

62. Guatteo R, Beaudeau F, Joly A, Seegers H. *Coxiella burnetii* shedding by dairy cows. Vet Res. 2007;38:849–60.

63. Sutherland WJ, Freckleton RP, Godfray HCJ, Beissinger SR, Benton T, Cameron DD, et al. Identification of 100 fundamental ecological questions. J Ecol. 2013;101:58–67.

