## Supplementary material for "A generic multi-level stochastic modelling framework in computational epidemiology"

This document provides complementary information for the main article. S1 Appendix presents the multi-level agent classes used in the framework EMULSION to represent individuals or groups. S2 Appendix describes the alternative computation of bacteria quantities transported by wind. S3 Appendix provides additional figures. S4 Appendix presents the additional file provided with this supplementary material. Finally, S5 Appendix shows differences between model description files.

#### S1 Appendix Multi-level agent classes

The simulation architecture EMULSION's generic engine relies upon a multi-level agent-based wraping of individuals and organization levels, to wrap existing scales and paradigms.

All agents (subclasses of `EmulsionAgent`, cf. fig. S1 Fig) are situated in at least one `Environment` (the area where they live, perceive, and act). `AtomAgent` and its subclass `EvolvingAtom` represent individuals, the behavior of the latter being driven by at least one state machine.

Other classes are groups (subclasses of `GroupAgent`), which means they encapsulate an environment where other agents are located. Multi-level agent-based architectures can handle several kinds of groups, depending on the relations between the group and the underlying agents (Mathieu P., Morvan G., Picault S., 2018, "Multi-level agent-based simulations: Four design patterns". *Simul. Model. Pract. Theory*, in press). The subsequent groups are based on such typical situations.

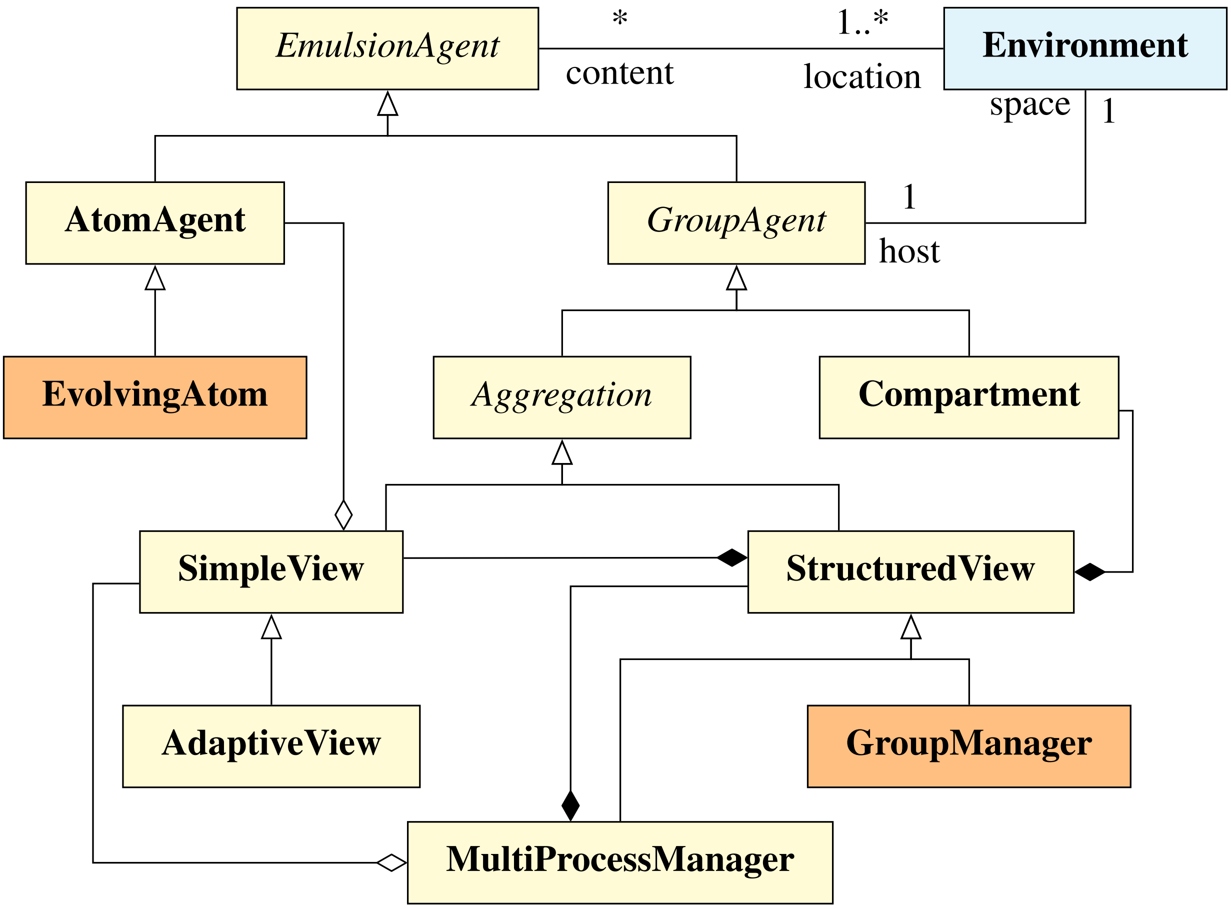

**S1 Fig:** UML class diagram for agents in the framework EMULSION. Agents with state-machine-driven behavior are identified with orange background. Triangular arrows denote specialization (also named "kind-of" relation), e.g. `EvolvingAtom` agents are a special kind of `AtomAgent`. Filled (resp. empty) diamonds represent the "composed of" (resp. "contains") relation, e.g. a `StructuredView` has to be composed of `SimpleView` agents, while a `SimpleView` can contain (but not necessarily) `AtomAgents`.

Among them, the simplest is the `Compartment`, suited when maintaining individual differences is not relevant. It simply aggregates homogeneous individuals into a quantity and can be tuned to act in a deterministic or stochastic way.

Conversely, `Aggregation` agents allow to gather individuals, according to key variables (e.g. health state, age, species...), in relation to a specific concern. Among them, `SimpleView` and its subclass, `AdaptiveView`, are intended to host atoms (individuals) and schedule their actions. The latter is also able to detect changes in individual variables to ensure grouping consistency.

`StructuredView` agents can host either `SimpleView` or `Compartment` agents, associated with possible values of concern-related variables. Finally, `GroupManager` agents are a kind of `StructuredView` endowed with a state machine, which determines how the state of agents changes over time.

`MultiProcessManager` agents are designed to control several concerns in the same simulation. Therefore, they are built using at least a `SimpleView` to hold atoms, and one or more `StructuredView` or `GroupManager` to handle each process involved in the model.

#### S2 Appendix Airborne transmission

The calculus of the quantity of bacteria $E_{\text{total}}$ present in the environment at instant $t$ for herd $i$ can be expressed as follows:

$$E^t_{i_{\text{total}}} = E^t_{i_{\text{local}}} + E^t_{i_{\text{aero}}} \tag{1}$$

where $E_{i_{\text{local}}}$ denotes the quantity of bacteria resulting from excretion of herd $i$ (handled by excretion actions in the health and lifecycle state machines, cf. figures S2 Fig–S5 Fig), and $E_{i_{\text{aero}}}$ the quantity received from each herd in the neighborhood:

$$E^t_{i_{\text{aero}}} = \sum_{j \neq i} E^t_{ij_{\text{plume}}} \quad (2)$$

$$\text{with:} \quad E^t_{ij_{\text{plume}}} = \zeta \times C(Q^t_j, x_{ij}, y_{ij}, z_{ij}) \times area \times height \quad (3)$$

$$\text{and:} \quad Q^t_j = \kappa \times \mu \times E^{t-1}_{j_{\text{local}}} \quad (4)$$

In the expressions above $Q^t_j$ represents the bacteria emission rate by herd $j$, with $\kappa$: proportion of bacteria eliminated from the within-herd environment becoming source for plume generation of herd $i$; $\mu$: elimination rate of bacteria; $\zeta$: contact rate between local animals and airborne bacteria; $area$: average surface occupied by a cow in the herd; $height$: average height where cows inhalate bacteria in the air.

The initial calculation of function $C$ was following a dispersion function (called in the article "Ermak-Stockie function") adapted from (Stockie, 2011 § 3.6), accounting for settling and deposition due to gravity:

$$\begin{aligned} C(Q, x, y, z) =& \frac{Q}{2\pi U \sigma_y \sigma_z} \exp(-\frac{y^2}{2\sigma_y^2}) \exp(-\frac{W_{set}(z-H)}{2K_z} - \frac{W_{set}^2 \sigma_z^2}{8K_z^2}) \\ & \times [\exp(-\frac{(z-H)^2}{2\sigma_z^2}) + \exp(-\frac{(z+H)^2}{2\sigma_z^2}) \\ & - \sqrt{2\pi}\frac{W_o \sigma_z}{K_z} \exp(\frac{W_o(z+H)}{K_z} + \frac{W_o^2 \sigma_z^2}{2K_z^2}) \text{erfc}(\frac{W_o \sigma_z}{\sqrt{2}K_z} + \frac{z+H}{\sqrt{2}\sigma_z})] \end{aligned} \quad (5)$$

In this function, $U$ denotes the mean wind velocity at time $t$; $H$, the altitude of the source herd; $W_{set}$ and $W_{dep}$, the settling and deposition velocities respectively; $K$, the diffusivity; erfc($x$), the complementary error function; $\sigma_y$ and $\sigma_z$, the standard deviations for dispersion coefficients corresponding to the atmospheric stability class C (3–5 m/s wind velocity, slightly unstable), as described in (Stockie, 2011); and $W_o := W_{dep} - \frac{1}{2}W_{set}$.

Since this approach only accounted for deposition of bacteria on the ground, we also tested the assumption that bacteria in the air were inhalated by cows. This is modelled by the much simpler "Gaussian plume" solution adapted from (Stockie, 2011 § 3.1):

$$\text{and:} \quad C(Q, x, y, z) = \frac{Q}{2\pi U \sigma_y \sigma_z} e^{-\frac{y^2}{2\sigma_y{}^2}} [e^{-\frac{(z-H)^2}{2\sigma_z{}^2}} + e^{-\frac{(z+H)^2}{2\sigma_z{}^2}}] \quad (6)$$

Each function was implemented as a specific code add-on within EMULSION. The comparison between Ermak-Stockie function and Gaussian function is shown on figure 5 in the article.

**Reference:** Stockie JM. 2011 The Mathematics of Atmospheric Dispersion Modeling. *SIAM Rev.* 53, 349–372. (doi:10.1137/10080991X)

### S3 Appendix Additional figures

All figures below are related to the Q fever study presented in section "Results" of the main paper. **The role and value of all parameters are described in Supplementary file `Additional File 2.yaml`.**

**Reference:** Courcoul A, Monod H, Nielen M, Klinkenberg D, Hogerwerf L, Beaudeau F, et al. Modelling the effect of heterogeneity of shedding on the within herd *Coxiella burnetii* spread and identification of key parameters by sensitivity analysis. *J Theor Biol.* 2011;284:130–141. (10.1016/j.jtbi.2011.06.017)

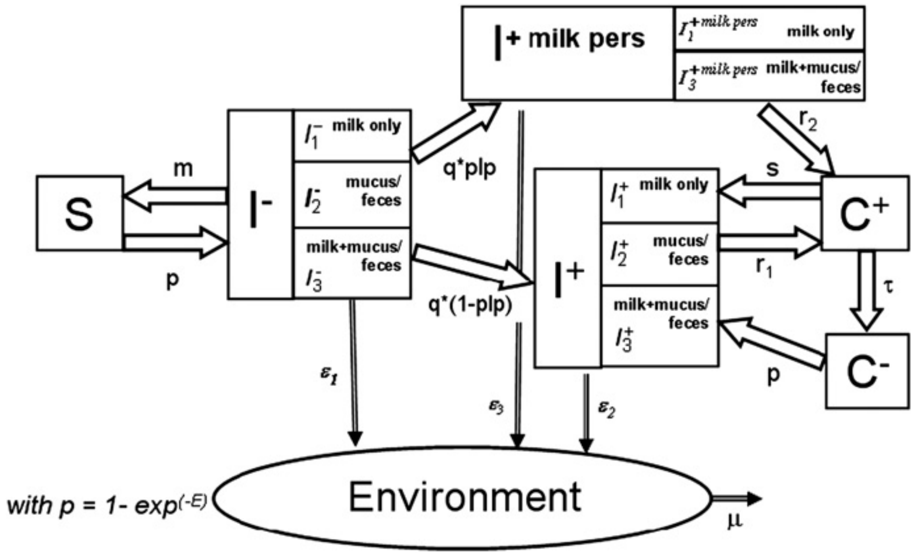

**S2 Fig:** Original flow diagram for the model of Q fever spread within a cattle herd, from Courcoul *et al.*, 2011.

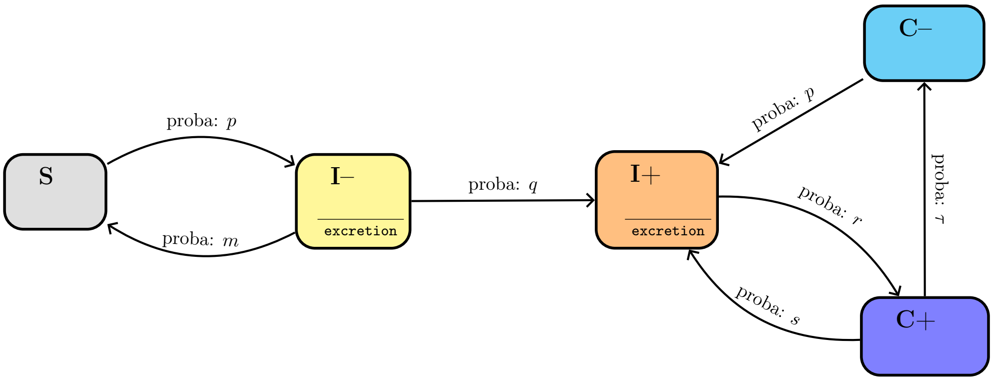

**S3 Fig:** State machine for the animal infection process in the initial ("simplified") model. Five health states are considered: Susceptible (S), Infectious without (I-) or with (I+) antibodies, Carrier i.e. non-infectious with (C+) or without (C-) antibodies. Specific actions (excretion of bacteria in the environment) are associated with infectious states. Since excretion has been reported to be intermittent, I- animals can become susceptible again.

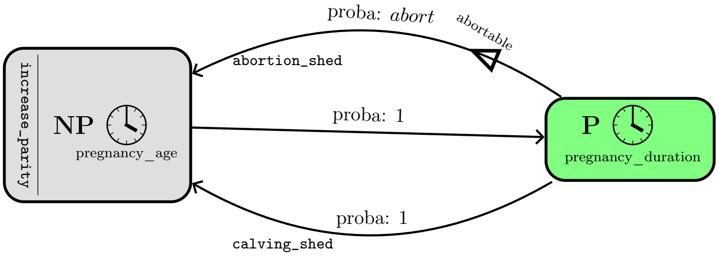

**S4 Fig:** State machine for the animal life cycle process in the initial ("simplified") model. Two states are considered: Pregnant (P) and Non-Pregnant (NP). Pregnant animals can return to Non-Pregnant either through calving after the pregnancy duration, of sooner through abortion (using the escape condition). Calvings and abortions are associated with a bacterial shedding peak materialized by an action.

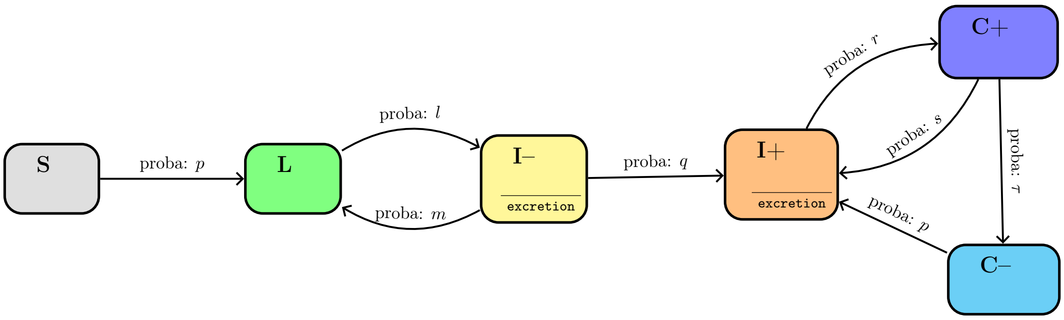

**S5 Fig:** Animal infection process after introducing a Latent state (L). Infectious animals cannot return to Susceptible state anymore, but only to the Latent state.

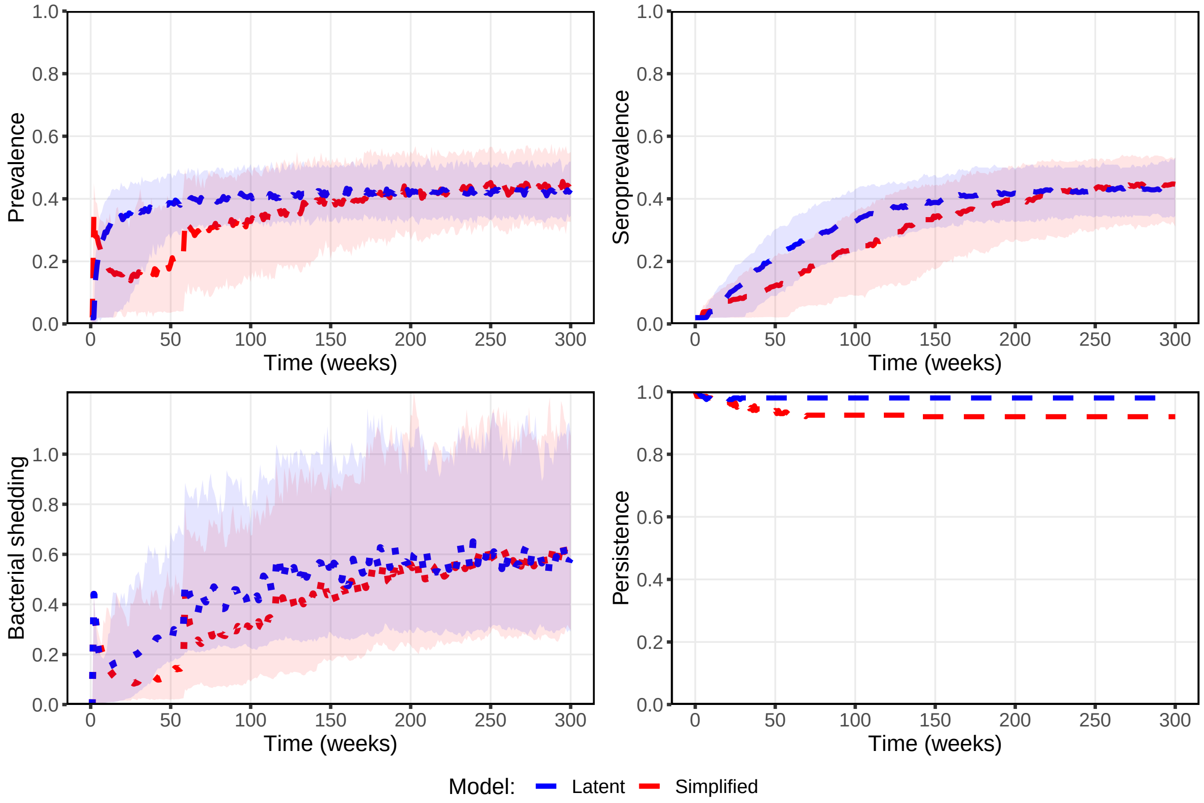

**S6 Fig:** Average (200 runs) and 10-90 percentiles for the prevalence (top left), seroprevalence (top right), pathogen shedding in environment (bottom left) and persistence (bottom right) comparing the model without (red) and with (blue) the Latent state. Results obtained after introducing a I+ animal just before calving in a fully susceptible herd.

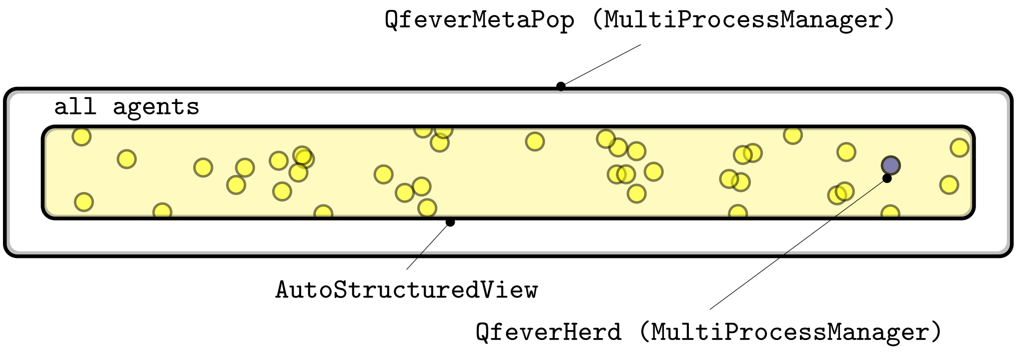

**S7 Fig:** Structure of the between-herd metapopulation model of Q fever. Each atom is actually a herd agent.

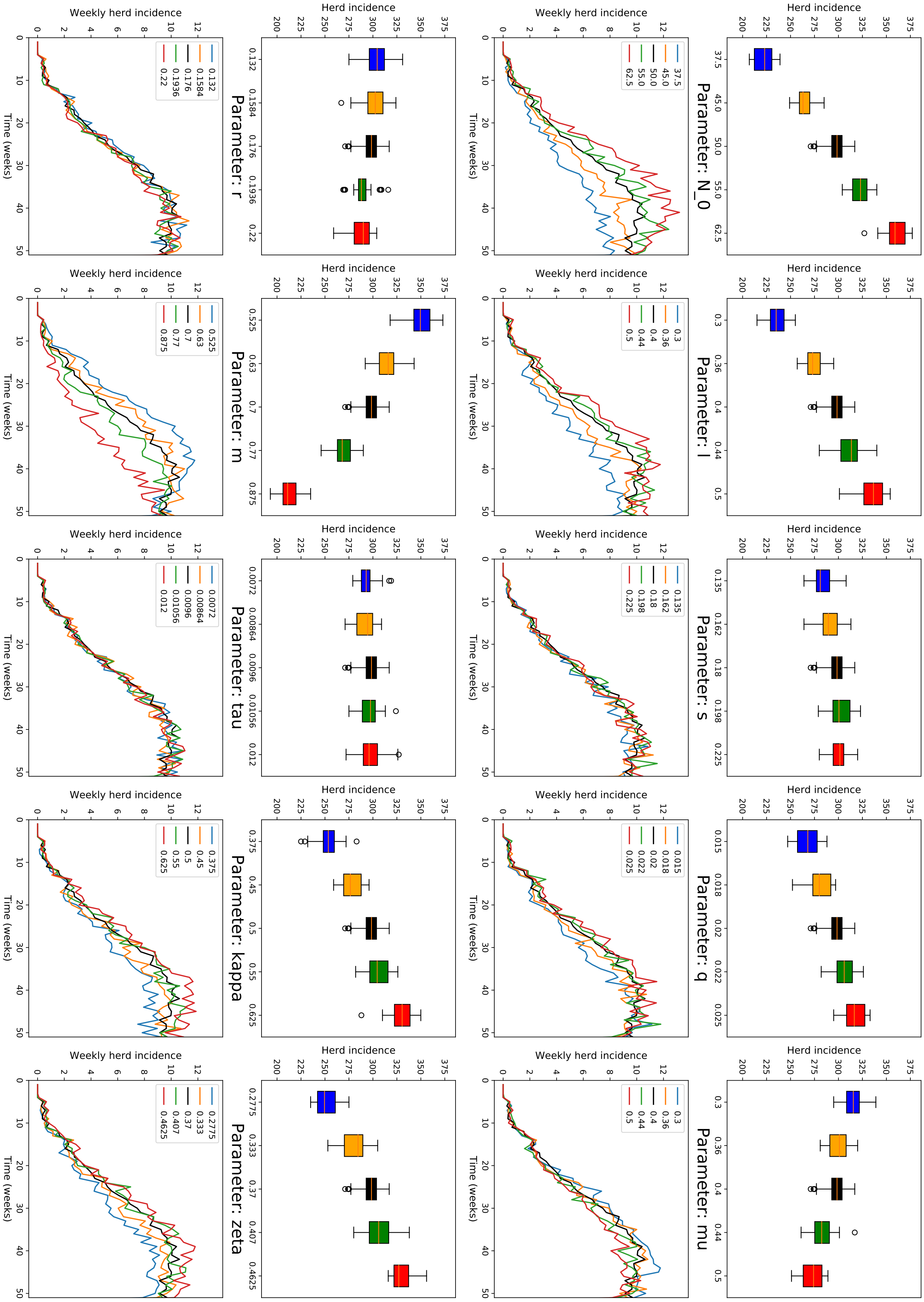

**S8 Fig:** Univariate sensitivity analysis (between-herd Q fever Latent state model) with respect to the main parameters. For each parameter: (top) distribution of herd incidences (seroprevalent herds) on 50 runs; (bottom) average weekly herd incidence (on 50 runs) over time. $N_0$: reference herd size; $l, s, q, r, m, \tau$: transition probabilities between health states (fig. S5 Fig); $\mu$: elimination rate of bacteria; $\kappa$: proportion of bacteria leaving local environment to contribute to aerial dispersion; $\zeta$: contact rate between local animals and airborne bacteria (reference values in black).

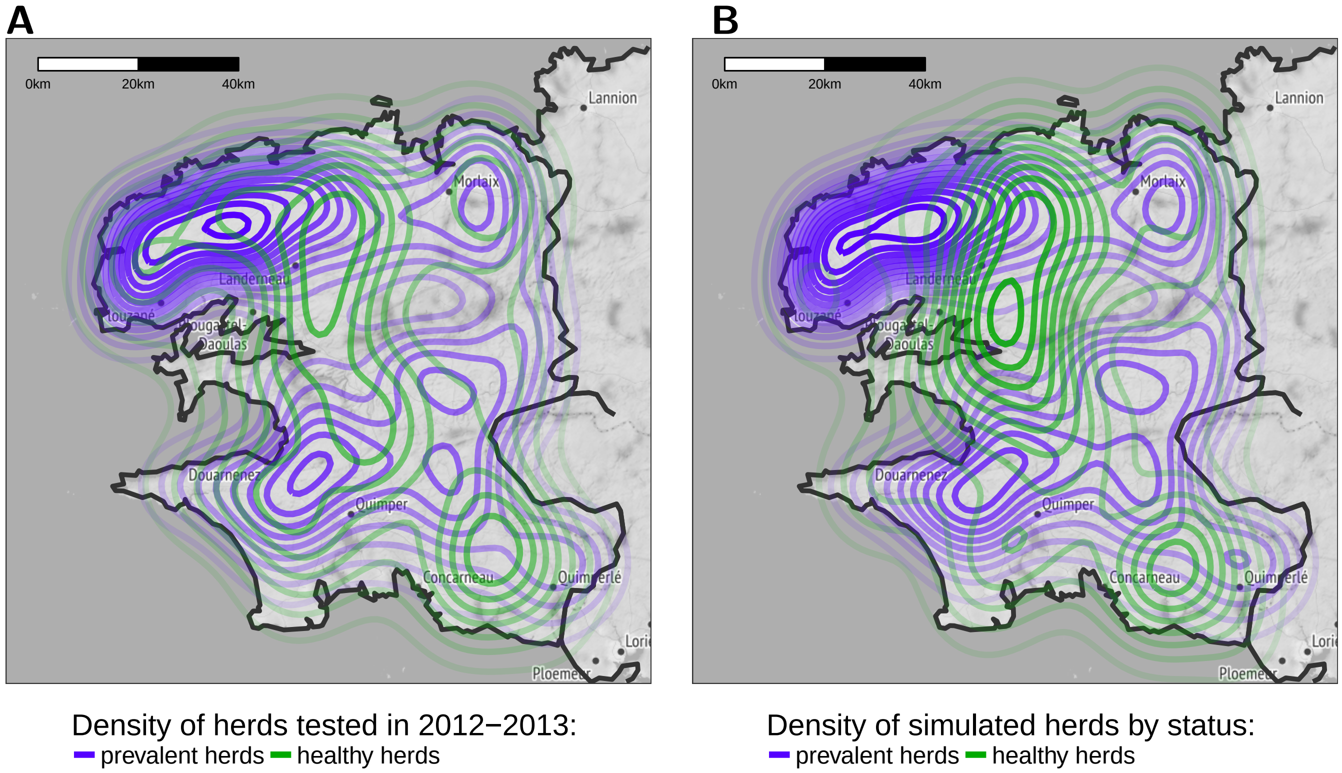

**S9 Fig:** Comparison of herd densities by seroprevalence (2D kernel density estimation). **(A)** Density of herds by seroprevalence status in 2012-2013 data. Prevalent herds aggregate herds tested seropositive either in 2012 or 2013, while healthy herds were seronegative both in 2012 and 2013. **(B)** Density of herds by health status in the simulations (50 runs). Herds were considered prevalent when they were reported seroprevalent in 2012 data, or if they were incident in a number of runs above the median number of incident runs; otherwise, they were considered healthy.

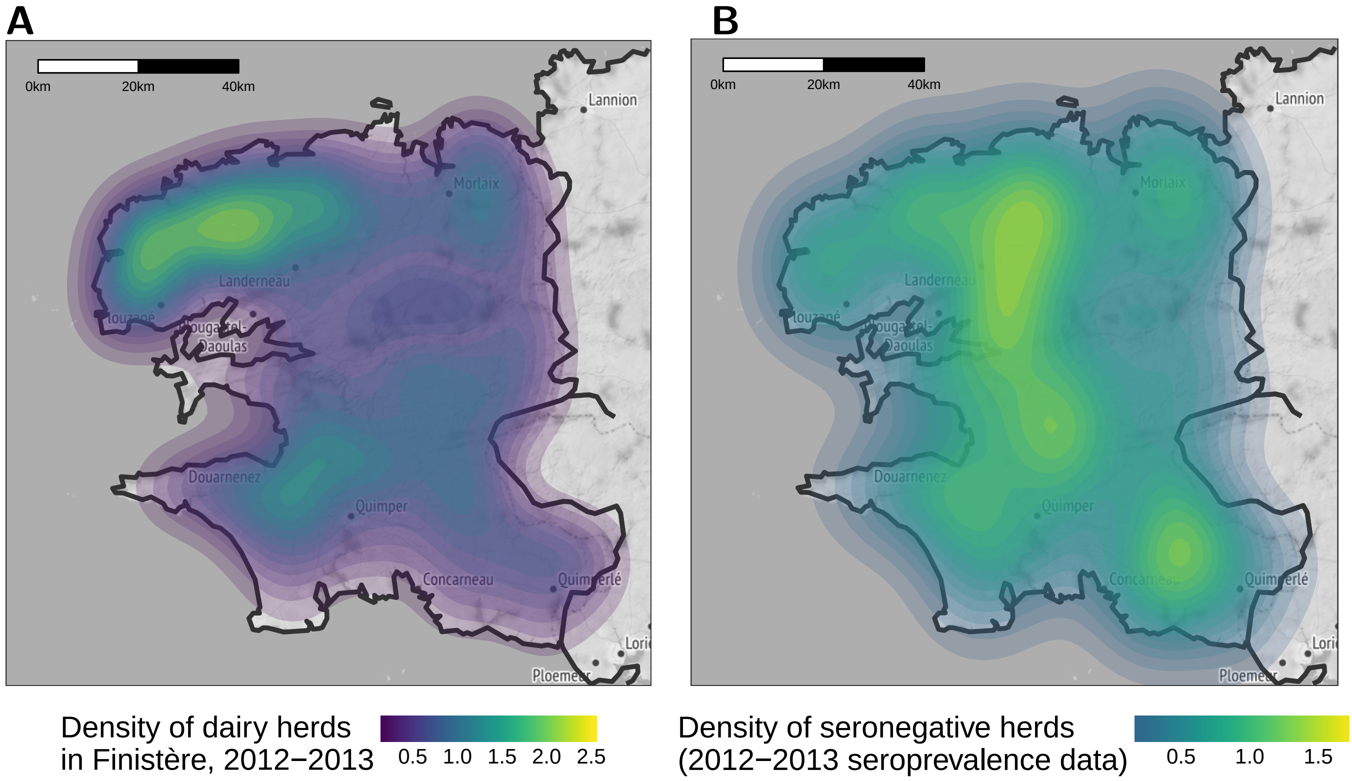

**S10 Fig:** **(A)** Density of dairy herds in Finistère for the study period (2012-2013). **(B)** Density of herds tested seronegative in 2012 and 2013.

#### S4 Appendix Files

Supplementary file below contains the YAML configuration for describing the simplified within-herd model of Q Fever (specifications for YAML data serialization standard are available at: `http://yaml.org`). This file is structured hierarchically, based on nested key-value pairs and lists. We provide below the whole listing of this file, colorized for better readability.

```yaml
---
# NAME OF THE MODEL
model_name: QFever

# INFORMATION ON THE DISEASE
info:
  # associated pathogen
  pathogen: Coxiella burnetii

# TIME INFORMATION
time_info:
  # time unit used for specifying parameter values
  time_unit: weeks
  # simulation time step, expressed in time units
  delta_t: 1
  # date where simulation starts
  origin: 'January 1, 2012'

# LEVEL OF ENTITIES INVOLVED IN THE SIMULATION
levels:
  individuals:
    super:
      module: emulsion.agent.atoms
      class_name: AtomAgent
    module: emulsion.examples.qfever.simplified_agents
    class_name: Cow
    description: 'class for representing individuals (here, adult cows)'
  herd:
    super:
      module: emulsion.agent.managers
      class_name: MultiProcessManager
      master:
        module: emulsion.agent.views
        class_name: SimpleView
    module: emulsion.examples.qfever.simplified_agents
    class_name: QfeverHerd
    description: 'class for representing herds'

# LIST OF PROCESSES OCCURRING AT EACH LEVEL (IN ORDER)
# processes which are not handled by a State Machine are expected to
# be methods within the reference level class (e.g. QFeverHerd)
processes:
  herd:
    - culling_process
    - renewal_process
    - infection
    - lifecycle
```

```
    - parity_grouping
    - bacterial_dispersion

# DESCRIPTION OF GROUPINGS. Agents from one level can be dynamically
# gathered, according to specific variables. Processes which are not
# based on specific methods in Python classes rely on groups
# associated with state machines
compartment_description:
  herd:
    parity_grouping:
    # The parity_grouping process is a simple grouping on individuals
    # based on the 'parity' statevar
      key_variables: [parity]
      compart_manager:
        # Agent class in charge of managing Parity groups
        module: emulsion.examples.qfever.simplified_agents
        class_name: ParityDictComp
      compart_class:
        # agent class which holds invidivuals with the same parity
        module: emulsion.agent.views
        class_name: AdaptiveView
    lifecycle:
    # The lifecycle process is based on a state machine and the
    # corresponding variable
      # The machine name is also the name of the variable
      # which holds the current state of the agent
      machine_name: life_cycle
      key_variables: [life_cycle]
      compart_manager:
        module: emulsion.agent.managers
        class_name: GroupManager
      compart_class:
        module: emulsion.examples.qfever.simplified_agents
        class_name: ObserverLifeStateComp
    # The infection process is based on a state machine and the
    # corresponding variable
    infection:
      machine_name: health_state
      key_variables: [health_state]
      compart_manager:
        module: emulsion.examples.qfever.simplified_agents
        class_name: HealthStateDictComp
      compart_class:
        module: emulsion.examples.qfever.simplified_agents
        class_name: ObserverHealthStateComp

# DESCRIPTION OF THE STATE MACHINES involved in the model
state_machines:
  life_cycle:
  # DESCRIPTION OF THE STATE MACHINE DRIVING COWS LIFE CYCLE
    # LIST of the states with their description
    states:
```

```yaml
    - NP:
        name: NotPregnant
        desc: 'Not in pregnancy period'
        fillcolor: lightgray
        on_enter:
          - action: increase_parity
        duration: pregnancy_age
    - P:
        name: Pregnant
        desc: 'pregnancy period'
        fillcolor: limegreen
        duration: pregnancy_duration
  # TRANSITIONS BETWEEN EACH HEALTH STATE
  # list of edges {from: source, to: dest, proba: probability of transition}
  transitions:
    - from: NP
      to: P
      proba: 1
    - from: P
      to: NP
      proba: 1
      on_cross:
        # action performed at calving (normal transition P -> NP)
        - action: calving_shed
          l_params: ['Q_calving']
    - from: P
      to: NP
      unless: 'abortable'
      proba: abort
      on_cross:
        # action performed when abortion occurs (premature return P -> NP)
        - action: abortion_shed
          l_params: ['early_abortion*Q_early_abort +
                     (1-early_abortion)*Q_late_abort']
health_state:
# DESCRIPTION OF THE STATE MACHINE DRIVING HEALTH STATES
  states:
    - S:
        name: Susceptible
        desc: 'non-shedder cows without antibodies'
    - I-:
        name: Infectious minus
        desc: 'shedder cows without antibodies'
        fillcolor: yellow
        on_stay:
          - increase: Etemp
            rate: 'Q_I_minus'
    - I+:
        name: Infectious plus
        desc: 'shedder cows with antibodies'
        fillcolor: orange
        on_stay:
          - increase: Etemp
```

```
              rate: 'Q_I_plus'
      - C+:
          name: Carrier plus
          desc: 'non-shedder cows with antibodies'
          fillcolor: royalblue
      - C-:
          name: Carrier min
          desc: 'non-shedder cows without antibodies, which were
          infected and had antibodies in the past'
          fillcolor: lightskyblue

    transitions:
      - from: S
        to: I-
        proba: p
        on_cross:
          - action: reset_last_infection
          - action: update_incidence_shedder
      - {from: I-, to: S, proba: m}
      - from: I-
        to: I+
        proba: 'q'
        on_cross:
          - action: update_incidence_sero
      - {from: I+, to: C+, proba: 'r'}
      - from: C+
        to: I+
        proba: s
        on_cross:
          - action: reset_last_infection
          - action: update_incidence_shedder
      - {from: C+, to: C-, proba: tau}
      - from: C-
        to: I+
        proba: p
        on_cross:
          - action: reset_last_infection
          - action: update_incidence_sero
          - action: update_incidence_shedder

# VARIABLES owned by agents
statevars:
  Etemp:
    desc: 'temporary bacteria excreted by the herd in its local
    environment within a time step'
  Eexcr:
    desc: 'total bacteria excreted by the herd in its local
    environment (cumulative)'
  Eplume:
    desc: 'total bacteria transported to the herd by neighboring herds
    (Plume model)'
  Eaero:
    desc: 'total bacteria transported by wind, which are inhalated by animals'
```

```yaml
nb_abortion:
  desc: 'number of abortions that occur in herd during the simulation'
nb_recursion_abortion:
  desc: 'number of abortions in herd that occur to cows which already
  aborted, throughout the simulation'
herd_population:
  desc: 'current population of herd (total size)'
abortable:
  desc: '1 if the animal is likely to abort, 0 otherwise. According
  to the model, an animal could abort during the 3 weeks following
  infection or resumption of shedding. Abortion could occur at any
  time of the gestation.'
early_abortion:
  desc: '1 if abortion occurs early, 0 otherwise'
last_infection:
  desc: 'number of weeks elapsed since last infection occured'
origin_herd_id:
  desc: 'origin herd id when a cow is created'

# HERD DEMOGRAPHY VARIABLES
# Pass by kwargs when initialization
# Values are calculated from detention data (BDNI)
init_pop:
  desc: 'herd size at the beginning of the simulation'
annual_renew_prop:
  desc: 'annual proportion of renewal process (herd size)'
culling_proba_0:
  desc: 'Culling probability corresponding to parity 0'
culling_proba_1:
  desc: 'Culling probability corresponding to parity 1'
culling_proba_2:
  desc: 'Culling probability corresponding to parity 2'
culling_proba_3:
  desc: 'Culling probability corresponding to parity 3'
culling_proba_4:
  desc: 'Culling probability corresponding to parity 4'
culling_proba_5:
  desc: 'Culling probability corresponding to parity 5'
culling_proba_6:
  desc: 'Culling probability corresponding to parity 6 and more'
dist_parity_0:
  desc: 'Composition proportion of parity 0 cow in herd'
dist_parity_1:
  desc: 'Composition proportion of parity 1 cow in herd'
dist_parity_2:
  desc: 'Composition proportion of parity 2 cow in herd'
dist_parity_3:
  desc: 'Composition proportion of parity 3 cow in herd'
dist_parity_4:
  desc: 'Composition proportion of parity 4 cow in herd'
dist_parity_5:
  desc: 'Composition proportion of parity 5 cow in herd'
```

```
# MODEL PARAMETERS AND EXPRESSIONS
parameters:
  pregnancy_age:
    desc: 'time elapsed since last calving, before pregnancy starts'
    value: 15
  pregnancy_duration:
    desc: 'duration of the pregnancy period'
    value: 40
  abort:
    desc: 'abortion probability'
    value: '1-(1-0.02)^(1/3)'
  max_abort:
    desc: 'maximum pregnancy week when abortion can occur'
    value: 30
  early_abortion_date:
    desc: 'number of weeks below which abortions are considered early'
    value: 15
  Etotal:
    desc: 'total bacteria deposited in the environment'
    value: 'Eexcr + Eaero'
  p:
    desc: 'infection probability'
    value: '1-exp(-Etotal*N_0/herd_population)'
  N_0:
    desc: 'reference population (for normalization)'
    value: 50
  m:
    desc: 'transition probability I- => S'
    value: 0.7
  l:
    desc: 'transition probability L => I-'
    value: 0.4
  q:
    desc: 'transition probability I- => I+'
    value: 0.02
  r:
    desc: 'transition probability I- => C+'
    value: 0.176
  s:
    desc: 'transition probability C+ => I+'
    value: 0.18
    source: 'A. Courcoul 2010'
  tau:
    desc: 'transition probability C+ => C-'
    value: 0.0096
  Q_I_minus:
    desc: 'average shedding quantity for cows in I- state'
    value: 0.003
    source: 'Courcoul 2010'
  Q_I_plus:
    desc: 'average shedding quantity for cows in I+'
    value: 0.004
    source: 'Courcoul 2010'
```

```
Q_calving:
  desc: 'quantity shedded at calving'
  value: 0.5758120438
  source: 'Average in Courcoul 2011'
Q_early_abort:
  desc: 'quantity shedded at early abortion'
  value: 0.013066666666666667
  source: 'Average in Courcoul 2011'
Q_late_abort:
  desc: 'quantity sheded at late abortion'
  value: 0.392
  source: 'Average in Courcoul 2011'
mu:
  desc: 'elimination rate of C. burnetii'
  value: 0.4
  source: 'Courcoul 2010 0.17 - 0.24'
kappa:
  desc: 'proportion of bacteria eliminated from local environment
  that contribute to plume dispersion'
  value: 0.5
zeta:
  desc: 'contact rate between local cows and bacteria transported by plume'
  value: 0.28
culling_threshold:
  desc: 'threshold for culling process'
  value: 0.85
renewal_threshold:
  desc: 'threshold for renewal process'
  value: 1.15
renew_proba:
  desc: 'probability of buying a new heifer per week per processed cow'
  value: '1-(1-annual_renew_prop)^(1/52)'
neighborhood_distance:
  desc: 'maximum distance (m) for takin into account wind transportation'
  value: 10000
surface_per_cow:
  desc: 'average surface (m^2) occupied per cow in a herd'
  value: 17
breath_height:
  desc: 'height (m) where cows breath'
  value: 2
depot:
  desc: '1 to activate the deposition model, 0 otherwise'
  value: 0
aero:
  desc: '1 to activate the inhalation model, 0 otherwise'
  value: 1
metapop_risk:
  desc: '1 to set the risk of buying an infected animal from outside
  of the metapop (external risk) to the average risk level of the metapopulation, 0
  to set the external risk to 0'
  value: 1
```

```
# ACTIONS that can be associated to states or transitions
actions:
  reset_cycle:
    desc: 'set the cycle week to 0 if nothing special occurs. In case
    of abortion, set the cycle week to a value depending on the week when
    abortion occurs'
  increase_parity:
    desc: 'increase the parity (number of calvings) of the animal'
  increase_cycle:
    desc: 'increase the number of weeks spent in each life cycle, and
    checks that it is strictly below the max_duration value associated
    to this state'
  abortion_shed:
    desc: 'quantity of bacteria shedded by a cow when an abortion occurs,
    the amount depends on the statevar early_abortion'
  calving_shed:
    desc: 'quantity of bacteria shedded by a cow when calving'
  reset_last_infection:
    desc: 'Reset the last_infection value when crossing S => I-, C+ =>
    I+ and C- => I+'
  update_incidence_shedder:
    desc: 'Update the herd incidence without antibodies'
  update_incidence_sero:
    desc: 'Update the herd incidence with antibodies'

# PERIODICITY OF OUTPUTS
outputs:
  herd:
    period: 1
  metapop:
    period: 51

# INITIALIZATION PARAMETERS
renew_animal:
  health_state: S
  life_state: NP
  parity: 0
  vaccinated: 0

init_susceptible_animal:
  health_state: S
  life_state: distribution
  parity: distribution
  vaccinated: 0

init_infected_animal:
  health_state: I+
  life_state: P
  _time_spent_life_cycle: duration_P
  parity: 0
  vaccinated: 0
```

```
# PREFERRED MODULES for symbolic computing
modules:
  - numpy
  - numpy.random
  - math
```

#### S5 Appendix Text

**S1 Text:** Modifications to do in the YAML configuration file describing the Q Fever model to move from within-herd to between-herd model. Lines highlighted in green starting with '+' must be added in the proper place. The `levels` section provides the description of the organization levels involved in the simulation (here, individuals and herds are already defined, and we want to add the metapopulation level). In the current state of EMULSION, an explicit identification of the agent classes required to build the level are required (this will be automatized in future versions): here, we create a class `QFeverMetaPop` based on `MultiProcessManager`, using an `AutoStructuredView` to hold all herds indexed by their ID. The `processes` section provides a list of processes occurring in each organization level (for the metapopulation: trade exchange and airborne dispersion of bacteria).

```
 ...
 levels:
   individuals:
     ...
   herd:
     ...
+  metapop:
+    super:
+      module: emulsion.agent.managers
+      class_name: MultiProcessManager
+      master:
+        module: emulsion.agent.views
+        class_name: AutoStructuredView
+        options:
+          key_variable: herd_id
+    module: emulsion.examples.qfever.simplified_agents
+    class_name: QfeverMetaPop
+    description: 'class representing a population of herds at regional scale'
 ...
 processes:
   herd:
     ...
+  metapop:
+    - wind_propagation
+    - exchange_animals
+    - outbox_to_inbox
+    - increase_step
 ...
```

**S2 Text:** Modifications to do in the YAML configuration file describing the Q Fever model to introduce the Latent (L) health state. Lines highlighted in green starting with '+' must be added in the proper place, while lines highlighted in red starting with '-' must be removed.

```
...
 state_machines:
   ...
   - health_states:
     states:
       - S:
           name: Susceptible
           desc: 'non-shedder cows without antibodies'
+      - L:
+          name: Latent
+          desc: 'latent cows (non-shedder and no antibodies)'
+          fillcolor: green
       - I-:
           name: Infectious minus
           desc: 'shedder cows without antibodies'
     ...
     transitions:
-      - from: S
+      - from: L
         to: I-
-        proba: p
+        proba: l
         on_cross:
           - action: reset_last_infection
           - action: update_incidence_shedder
-      - {from: I-, to: S, proba: m}
+      - {from: I-, to: L, proba: m}
+      - from: S
+        to: L
+        proba: p
       - from: I-
         to: I+
         proba: 'q'
 ...
```
